# Spectral methods for prediction uncertainty quantification in Systems Biology

**DOI:** 10.1101/2023.02.14.528500

**Authors:** Anna Deneer, Jaap Molenaar, Christian Fleck

## Abstract

Uncertainty is ubiquitous in biological systems. These uncertainties can be the result of lack of knowledge or due to a lack of appropriate data. Additionally, the natural variability of biological systems caused by intrinsic noise, e.g. in stochastic gene expression, leads to uncertainties. With the help of numerical simulations the impact of these uncertainties on the model predictions can be assessed, i.e. the impact of the propagation of uncertainty in model parameters on the model response can be quantified. Taking this into account is crucial when the models are used for experimental design, optimisation, or decision-making, as model uncertainty can have a significant effect on the accuracy of model predictions. We focus here on spectral methods to quantify prediction uncertainty based on a probabilistic framework. Such methods have a basis in, e.g., computational mathematics, engineering, physics, and fluid dynamics, and, to a lesser extent, systems biology. In this chapter, we highlight the advantages these methods can have for modelling purposes in systems biology and do so by providing a novel and intuitive scheme. By applying the scheme to an array of examples we show its power, especially in challenging situations where slow converge due to high-dimensionality, bifurcations, and spatial discontinuities play a role.

## 1 Introduction

Every mathematical model in systems biology is subject to uncertainty and incomplete knowledge [1–4]. This can be in the form of unknown model structure, unknown model parameters and imperfect experimental data. Characterizing and quantifying these sources is crucial, as the uncertainty can translate into inaccuracies in the model predictions. Information about the quality of model predictions is vital when applied as support for decision-making or optimization routines such as experimental design and parameter estimation [5]. The aim of uncertainty quantification (UQ) is to determine the likeliness of certain outcomes, given that some aspects of the system under study are not (exactly) known.

Generally, uncertainty is distinguished into two classes [6–8]. The first class is so-called aleatoric uncertainty. Aleatoric uncertainty stems from the intrinsic variability found in the system under consideration, for this reason it is also referred to as statistical uncertainty. For example, in the case of parameter estimation, this uncertainty is related to the fact that parameters may essentially vary over the system components (e.g., cells) [9], so that for the system as a whole only a distribution of parameter values can be estimated, and not one precise value per parameter. Gene expression noise is often a source of varying conditions (e.g., initial conditions or protein production rates) in cells resulting in variability of process parameters such as the protein production rate [9–13].

In contrast, the second class of uncertainty, termed epistemic (or systemic) uncertainty, is caused by a lack of information [6–8]. For example, in the case of parameter estimation, this may be caused by imperfect data sets that contain noisy, incoherent, or missing data points [14]. In such cases, the uncertainty can be reduced by performing extra experiments [15–17].

In biological systems both types of uncertainty are typically present [1]. In terms of modelling, both are usually dealt with by employing a probabilistic framework [18], in which model parameters can be represented according to a probability density function (PDF) [4, 6]. The choice of the type of PDF and the corresponding distribution of parameters is usually based on previous knowledge. For example, the case of a completely unknown parameter could be described by a uniform distribution on a broad (positive) interval. In other cases a parameter could be known to follow a normal or lognormal distribution with known mean and variance, established in previously performed experiments [19].

Among the field of UQ, Monte Carlo (MC) methods are most commonly used [20, 21]. In a MC approach the parameter PDFs are sampled and model responses for each sample recorded, thus providing a distribution of model outcomes and an indication of the uncertainty therein (e.g. by analyzing the distribution moments). These methods are simple in their implementation and are widely applicable. However, for models that have a large number of parameters or are computationally expensive, these MC procedures are often not feasible [6, 8].

As an alternative to MC, meta-modelling techniques are frequently adopted to deal with models that would otherwise be intractable. Support vector machines [22], artificial neural networks [23] and Bayesian networks [24] are examples of surrogate- or meta-modelling techniques used in systems biology. In this work, we focus on stochastic spectral methods, in particular polynomial chaos expansion (PCE), an approach that is widely used in engineering systems [7, 8] and to a lesser extent in biological systems [25–28] for UQ purposes. The aim of these approaches is to represent the model response as a series expansion.

The advantage of this representation is that an approximation of the model response is obtained for *all* values of the uncertainty parameters. This form allows immediate evaluation of statistics of the model outcome, either analytically or through sampling of the stochastic parameters, which can be done significantly faster than through MC methods for models that are problematic and computationally expensive [29].

This advantage comes at the cost of the need to calculate expansion coefficients. For this, two classes of spectral methods are in use. In the first class the governing equations of the model are reformulated such that each variable is represented by a spectral expansion. This results in a system of differential equations for the expansion coefficients and is known as intrusive spectral projection [6]. In the second class, consisting of so-called non-intrusive spectral projection approaches and followed in this paper, the expansion coefficients are determined using the model without changing the original model equations [30]. The advantage of this non-intrusive approach is that it requires only straightforward deterministic model evaluations and does not involve any reformulation of the model. It is particularly attractive in case of very large models where intrusive methods would become too laborious.

In the past, PCE has been shown to converge very slowly or not at all for models involving non-smooth functions [31, 32]. This is indeed a critical challenge for biological models, which often show complex, non-linear behaviour such as bifurcations and spatial discontinuities. In this work, we provide a scheme for non-intrusive spectral projection that may overcome these problems. It is easy to implement and we show its power through applying it to a number of biological models. In this scheme it is straightforward to use different basis functions for the expansion, such as Haar wavelets in case of models with bifurcations. We also present a simple segmentation method resulting in a piecewise continuous approximation of the model at hand which helps to deal with complex models avoiding the necessity of high order expansions. The examples treated in this paper each have a specific problem to overcome.

## 2 Methods

### 2.1 Spectral expansion

Let us consider a model **Ω** that depends on a vector of stochastic input parameters ***θ***. The model response *Y* can be any chosen quantity, e.g. the concentration of one of the model components or a function thereof. The uncertainty parameters *θ*_*l*_ are assumed to be independently distributed, each with PDF *P*_*l*_(*θ*_*l*_). So,

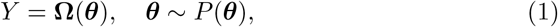

where *P* (***θ***) is the joint probability density function (PDF) for all *U* uncertainty parameters: 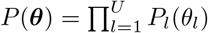. The case of correlated parameters is discussed in Sec. 2.2.3. For reasons of clarity, we restrict here the explanation to *Y* and *θ* being scalar functions. In the section Practical Aspects we show how to deal with more than one uncertainty parameter. Note that for correlated parameters the PDF would follow from the corresponding multivariate distribution.

The underlying model could be of any type, e.g., an ODE, a PDE, an algebraic, or a statistical model. This implies that *Y* may also depend on time and space. The challenge is to analyse the behaviour of *Y* as a function of ***θ***. In cases where the numerical evaluation of the underlying model takes a considerable amount of computational time this tends to obstruct any form of comprehensive analysis. In this paper we present the use of a method that aims at making this tractable. The idea is to replace the original model by a meta-model, which is achieved by representing the output *Y* in terms of an expansion. This meta-model can be constructed such that it represents the underlying model to a high degree of accuracy, with the advantage of being much faster to evaluate than the original model.

This meta-model can be used to determine the distribution of the model response *Y* or reconstruct the function accurately at given points in the parameter space. In the spectral approach the model response is represented by

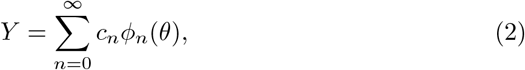

where *c*_*n*_ are the expansion coefficients (which can be time and/or space dependent) and *ϕ*_*n*_ are functions that are orthonormal with respect to the distribution of the uncertainty parameters as weight functions for an inner product, as dealt with in S1 Appendix A. For example, suitable basis functions for uniformly distributed parameters are Legendre polynomials, while for normally distributed parameters Hermite polynomials qualify. For any practical purpose the expansion needs to be truncated to a certain degree:

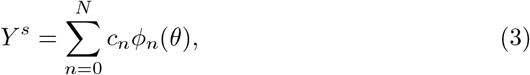

where *N* is the truncation degree. So, for such a meta-model *Y*^*s*^ *N* expansion coefficients have to be calculated. The advantage of (3) is that the statistics of the model response *Y* can be evaluated very fast, either analytically or through sampling of the parameters *θ*. The main computational cost of the expansion comes from the computation of the coefficients *c*_*n*_. Below, we provide an easy-to-implement scheme for the calculation of these coefficients.

We use a non-intrusive approach to the spectral expansion, i.e., we treat the model equations as a black box, not requiring any tailoring to the equations to include the parameter uncertainties. The most commonly used method for non-intrusively determining the coefficients is through Gaussian quadrature schemes [6]. Here, we propose an alternative scheme. It is applicable to any set of orthonormal functions, allowing the flexibility to tackle different modelling challenges. A key feature in the scheme is the introduction of the symmetric matrix:

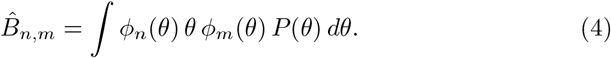

Its eigenvalues *λ*^(*l*)^, *l* = 1, 2, …, are real and its eigenvectors *u*^(*l*)^ orthonormal. In the appendix we show that these eigenvalues and eigenvectors can be used to derive an expression for the coefficients *c*_*n*_. After substitution, Eq. (3) then reads as

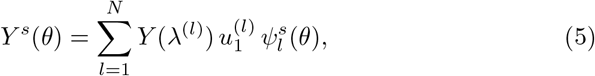

Where

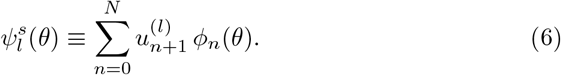

The striking point here is that this expansion requires to evaluate the model only *N* times, namely for each of the eigenvalues *λ*^(*l*)^, *l* = 1, …, *N* . Note that all terms in the expansion that do not depend on the uncertainty parameter *θ* can be calculated in advance, so once and for all. This saves computation time for any future application. For models that take a long time to evaluate the use of Eq. (5) is a very fast alternative, compared to e.g. a Monte-Carlo approach. What’s more, statistical moments like mean and variance follow directly from the expansion coefficients [6], thus requiring no further calculation. Note that expansion Eq. (5) is only exact in the limit *N*→ ∞ . Taking a finite value for *N* introduces an inaccuracy. Therefore, *N* must be chosen with care and it is often not obvious beforehand which value of *N* will give reliable results. We will showcase in the examples underneath how one may deal with the choice of *N*. In the subsection Segmentation in the next section, we propose a very simple adjusted scheme to deal with cases where a high degree might result in infeasible computational times.

### 2.2 Practical aspects

Here, we treat some specific aspects of the method presented above.

#### 2.2.1 PDFs and basis functions

We already mentioned Legendre and Hermite polynomials as typical basis functions for PCE. Legendre polynomials are defined over [ −1, 1] and are orthogonal with respect to the uniform distribution. The Hermite polynomials are defined over ℝ and are orthogonal with respect to the Gaussian distribution. Both polynomials can be normalized with appropriate prefactors to ensure orthonormality. These two families of polynomials are most commonly used to represent biological parameters. The uniform distribution is typically applied in uninformed cases and the lognormal distribution in cases where there is prior information available on a parameter. In practice, sometimes other classical orthogonal polynomial families are appropriate, e.g., Laguerre polynomials for Gamma distributions. Also, non-polynomial functions may be applied, such as spherical harmonics. The approach presented here can be used for any set of orthonormal functions.

The Legendre and Hermite polynomials are defined for standard uniform 𝒰 ( −1, 1) and standard normal 𝒩 (0, 1) variables, respectively. In practice, the biological parameters are often not restricted to the corresponding intervals. In these cases we have to apply an isoprobabilistic transformation. For example, to obtain a normally distributed random variable *k* with mean *µ* and variance *σ*, so *k* ∼ 𝒩 (*µ, σ*), from *θ* ∼ 𝒩 (0, 1), we need the transform

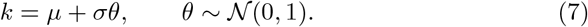

To obtain a uniformly distributed random variable *k* on the interval [*a, b*], so *k* ∼ 𝒰 (*a, b*), from *θ* ∼ 𝒰 (−1, 1), we need the transform

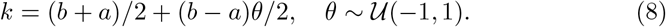

A lognormally distributed random variable *k* ∼ Lognormal(*µ, σ*) is obtained from *θ* ∼ 𝒩 (0, 1) via the transformation

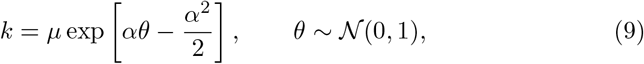

where 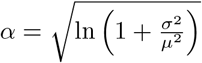.

#### 2.2.2 Multiple uncertainty parameters

Typically, biological models involve more than one random parameter, which means that the PCE basis {*ϕ*_*n*_(***θ***), *n*∈ℕ^*M*^} is multivariate. Extending (5) to the M-dimensional case is straightforward:

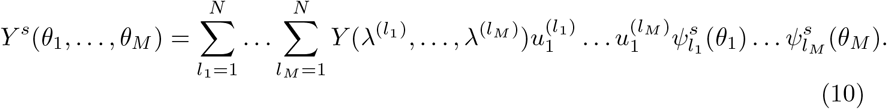

Similar to MC approaches, PCE suffers from the curse of dimensionality [7, 33]. Note from (10) that the number of times the model has to be evaluated scales as *N* ^*M*^, where *N* is the expansion order and *M* the number of uncertainty parameters.

#### 2.2.3 Correlated uncertainty parameters

To handle correlated parameters we consider the approach of transforming the basic functions. We discuss the case of two correlated parameters. In section S2 Supplementary Information we show the general case. Given that *P* (*θ*_1_, *θ*_2_) is the distribution for the uncertainty parameters *θ*_1_ and *θ*_2_, *P* (*θ*_1_) and *P* (*θ*_2_) are the corresponding marginalised distributions, and *P* (*θ*_1_, *θ*_2_) ≠ 0, ∀*θ*_1_, *θ*_2_, the new basis functions can be written as (for details see Supplementary Information S2):

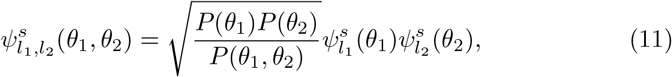

where the functions 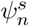 are orthonormal with respect to *P* (*θ*) and are given by Eq. 6. The expansion of a function depending on correlated parameters can be written as:

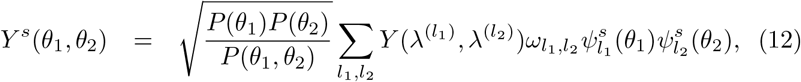

With

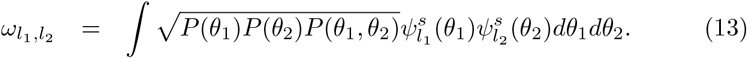

#### 2.2.4 Segmentation

In cases where an expansion to high order is needed to obtain the requested approximation accuracy or the model is computationally expensive, one needs to refer to optimised or effective methods [8, 14]. Models that show complex response surfaces (e.g., bifurcations) will require high order expansions to capture the inherent complexity. This is problematic since it not only requires to evaluate the model often, but also leads to time consuming summations in (10). To overcome these problems, we present here a simple and easy to implement scheme that segments the parameter intervals into subintervals, yielding a piecewise continuous approximation of the original function. Within each of these segments we then perform a separate expansion. In this approach we have to deal with a trade-off: the number of expansions is multiplied, but per expansion we have a (much) lower order of expansion. Below we argue why the second positive aspect greatly counterbalances the first negative aspect.

To determine the segments we define a scaling function *g*_*m*_ with *M* ∈ ℕ and *L* ∈ ℝ by:

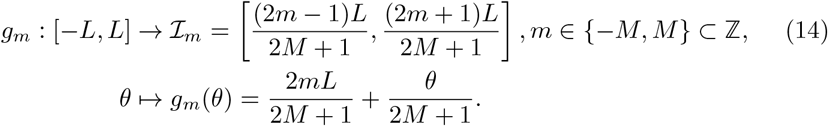

This scaling function *g*_*m*_ divides the interval [−*L, L*] into 2*M* + 1 segments ℐ_*m*_ of equal length. Of course, *L* must be larger than or equal to any value of *θ*. For example, consider the case *L* = 1. Then, the whole interval is [ −1, 1]. For a segmentation granularity of *M* = 1, this interval is divided into three subintervals: ℐ_*−*1_ = [−1, −0.33], ℐ_0_ = [−0.33, 0.33] and ℐ_+1_ = [0.33, 1]. The expansion of *Y* on any subinterval ℐ_*m*_ is given by:

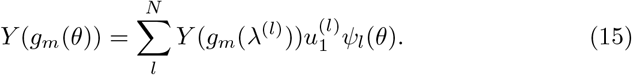

Upon a variable transformation *y* = *g*_*m*_(*θ*), Eq. 15 becomes:

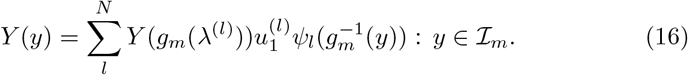

After segmentation, the expansion on the interval [ −*L, L*] as a whole is a superposition of the expansions on the subintervals:

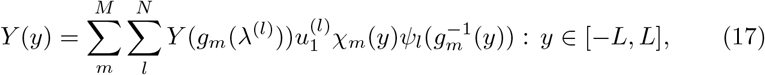

where *χ*_*m*_(*y*) is an indicator function for selecting the correct segment:

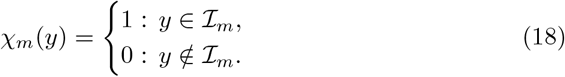

We can also define an index function to select *m*^∗^ ∈ {−*M, M*} for which *χ*_*m*_(*y*) =1:

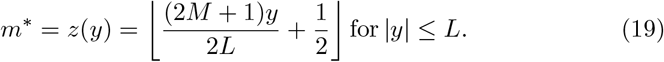

Using both index functions we can finally write the segmented reconstruction as:

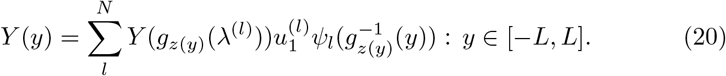

Segmentation allows the use of lower order polynomials while maintaining the same accuracy (assuming a sensible choice for *N* and *M*) as a non-segmented higher order expansion. The number of model solutions required now scales as (2*M* + 1)^*K*^*N* ^*K*^, where *K* is the number of uncertainty parameters, *M* the segmentation granularity, and *N* the expansion order. The reduced accuracy by using a lower order expansion is compensated for by evaluating the model more often, as a result of zooming in. Expanding up to the *N* -th order for *p* uncertainty dimensions requires solving the system *N* ^*p*^ times. Reconstruction requires the summation of *N* ^2*p*^ terms. Therefore, it is advantageous to keep *N* as low as possible. Normally, the reconstruction error is large for low *N*, but this is mitigated by segmentation. Segmentation requires to evaluate the system (2*M* +1)^*p*^*N* ^*p*^ times, but due to segmentation *N* can be taken much smaller.

To illustrate this with an example, we take a system with 2 species of interest and 5 uncertainty parameters *θ*_*i*_. The expansion order is taken as *N* = 8. This implies summing over 2 × 8^10^ = 2, 147, 483, 648 terms per time point and per parameter set (*θ*_1_, …, *θ*_5_). In the case of segmented expansion, we can choose a lower *N*, for example *N* = 3 with a segmentation granularity of *M* = 1. The number of terms to be summed over is 2×3^10^ = 118, 098. This is dramatically more efficient and stems from the fact that one only has to determine the segment in which the parameter set (*θ*_1_, .., *θ*_5_) falls and choose the corresponding expansion coefficients. In the Results section we will present Example II in which segmentation indeed proves to be very beneficial.

#### 2.2.5 Haar wavelet expansion

Traditional PCE methods are known to have difficulties with capturing discontinuous behaviour [31, 32]. Spectral convergence is only observed when solutions are sufficiently regular and continuous. Just like Fourier expansions, PCE suffers from Gibbs phenonema at discontinuities, resulting in slow convergence [8]. Haar wavelets have been suggested to overcome these difficulties [14, 32]. In contrast to global basis functions like the aforementioned polynomial systems, wavelet representations lead to localized decompositions, resulting in increased robustness at the cost of a slower convergence rate [8, 14]. Here, we discuss that Haar wavelets can be easily incorporated in the framework presented above and in Example IV underneath we show how they can be applied in practice.

As mother wavelet we take

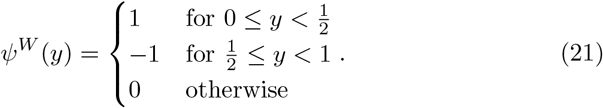

By introducing a scaling factor *j* and a sliding factor *k*, we may construct the wavelet family

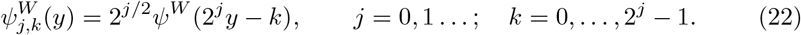

Given the uncertainty parameter *θ* with its cumulative distribution function *F* (*θ*), we define the basis functions as

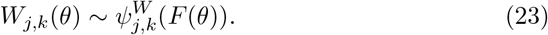

By concatenating the indices *j* and *k* into one index *i*≡2^*j*^ + *k*, we may expand the meta-model *Y* similarly as we did in Eq. (2):

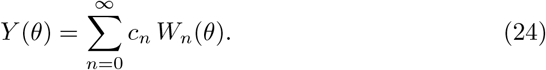

#### 2.2.6 Sensitivity analysis

In sensitivity analysis one quantifies the effects of changes in the parameters on the variability of the model response. Here, we show how our PCE approach allows for sensitivity analysis in an elegant way. In the case of local sensitivity analysis, small parameter variations around a certain point in parameter space are used to determine the effect on the model output [34]. This sensitivity is estimated via calculation of the partial derivatives of model output with respect to parameters, evaluated in that point [35]. Alternatively, global sensitivity approaches do not specify a specific point in parameter space [36]. For example, Sobol indices are a popular sensitivity measure as they provide a measure of global sensitivity and accurate information for most models [37]. Sobol indices are based on the decomposition of the variance of the output *Y* as a function of the contribution of the parameters (and possibly their combination), also called the ANalysis of VAriance, or ANOVA [37]. Thanks to the orthonormality of basis functions in PCE, Sobol indices can be determined analytically from the coefficients of the PCE [38, 39] So, once these coefficients are known, one gets the Sobol indices nearly for free. Given the PCE expansion of output *Y*, the total variance of the model output is given by

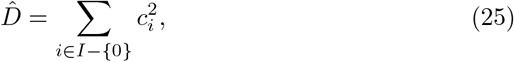

where I is the multi-index set of all variables and *c*_*i*_ the expansions coefficients. The 0^th^ coefficient is not included as this is a constant. The partial variance is given by

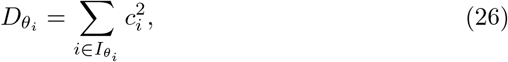

where 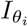 is the multi-index set of parameter *θ*_*i*_, i.e. where the *i*^th^ term in the multi-index is larger than 0. The Sobol indices are then given by

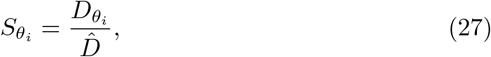

In this way the relative contribution of parameter *θ*_*i*_ to the variance of the output is easily calculated.

### 2.3 Monte Carlo sampling

In the examples shown in this paper, we have compared PCE with Monte Carlo sampling. In these cases, we have used Quasi Monte Carlo methods using Sobol sequences, which show a faster rate of convergence than standard sequences of pseudorandom numbers [40]. In order to test for convergence, we have used the so-called blocking method. In the blocking method the error is estimated in a straightforward manner. The quantity of interest (i.e. the model response *Y*) is divided into several groups (or blocks). Then, for each block, we determine the moment of interest (e.g. mean or variance). The spread (variance) of values between blocks gives an estimate of the error. Finally, to test for convergence, we choose a threshold on the between-block variance. Note that the convergence rate of MC methods do not depend on the dimensionality of the parameter space that is being sampled, unlike PCE. MC merely scales by 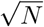, where *N* is the number of samples.

### 2.4 Summary of implementation

In this section we provide an overview of the steps needed to arrive at a metamodel using PCE:

1. Determine which of the model parameters may show stochastic behaviour and decide upon an appropriate PDF for such parameters.

2. Choose a truncation degree *N* .

3. Based on the PDF in the previous steps, calculate the appropriate basis functions *ϕ*_*n*_(*θ*), *n* = 0, 1, 2 …, *N* .

4. Determine the *N*× *N* matrix 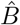 as defined in 4. For example, for Legendre polynomials 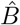 reads as

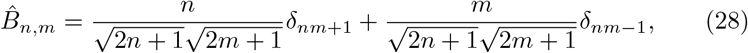

where *n, m* = 0, 1, 2 …, *N* .

For Hermitian polynomials 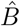 reads as:

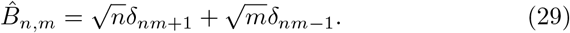

5. Calculate the eigenvalues *λ*^(*l*)^, *l* = 1, 2, …, *N* and orthonormal eigenvectors *u*^(*l*)^.

6. Calculate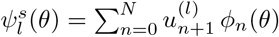 .

7. Calculate *Y* (*λ*^(*l*)^), *l* = 1, 2, …, *N* by evaluating the model *N* times.

8. Arrive at the metamodel 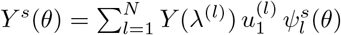.

9. Eventually, apply post-processing through, e.g., sensitivity analysis.

## 3 Results

To test the performance of the present PCE approach in biological simulations, we have chosen four typical examples. Through these examples, we show how to deal with several challenges usually encountered in systems biology.

The first example has only one uncertainty parameter. Its simplicity allows comparison between the results of our approach with an exact solution.

The second example concerns a biochemical reaction network and is higher dimensional, i.e., it contains more than one uncertainty parameters. We use it to highlight the advantages of segmentation.

The third example is the glycolytic oscillator, which shows bifurcations, i.e., different dynamic behaviour for different parameter sets [41]. We use it to demonstrate the power of global sensitivity analysis, which in the PCE framework can be achieved without significant additional computational costs once the PCE coefficients have been calculated. In addition, this example allows us to show the use of mixed expansions, since the parameter PDFs follow different distributions. This leads to a combination of different families of basis functions, thus highlighting the flexibility of the PCE approach when applied to varying input uncertainties.

The last two examples have a spatial dimension. First, we consider the Schnakenberg model which is a well-known model of pattern formation and comes with challenges such as shifts from non-patterning to patterning regions [42]. In this example we demonstrate the advantage of using Haar wavelets over polynomial basis functions for systems with bifurcations. Second, we study a model describing pattern formation in plants, more specifically patterning of the hairs found on top of leaves, so-called trichomes [43]. In this model we show how to adapt the approach such that the computational costs are reduced as much as possible by carefully choosing the quantity of interest, without changing the standard set of steps.

### 3.1 Example I. Exponential decay: comparing performance of PCE to MC and an analytical solution

For this example case, we consider the extremely simple reaction system consisting of one decaying species:

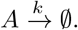

Its dynamics are described by *A*(*t*) = *A*_0_ exp^*−kt*^, where *A*_0_ is the initial concentration of *A* at *t* = 0 and *k* the rate of decay. We test the PCE method against:

*1)* the exact, analytical solution and *2)* the classical Monte Carlo approach.

The quantification of the sources of uncertainty constitute the second step in the analysis. This entails identification of the parameters that are unknown and modelling them in a probabilistic context. In this case, we assume that *k* is distributed according to a lognormal distribution with known mean and variance, i.e. *k*∼ Lognormal(*µ, σ*), and we choose *µ* = 0.5, *σ* = 0.2. The PDF for *k* is shown in Figure 1 and the derivation for the exact PDF for *A* is given in the Supplementary Information.

**Figure 1.**
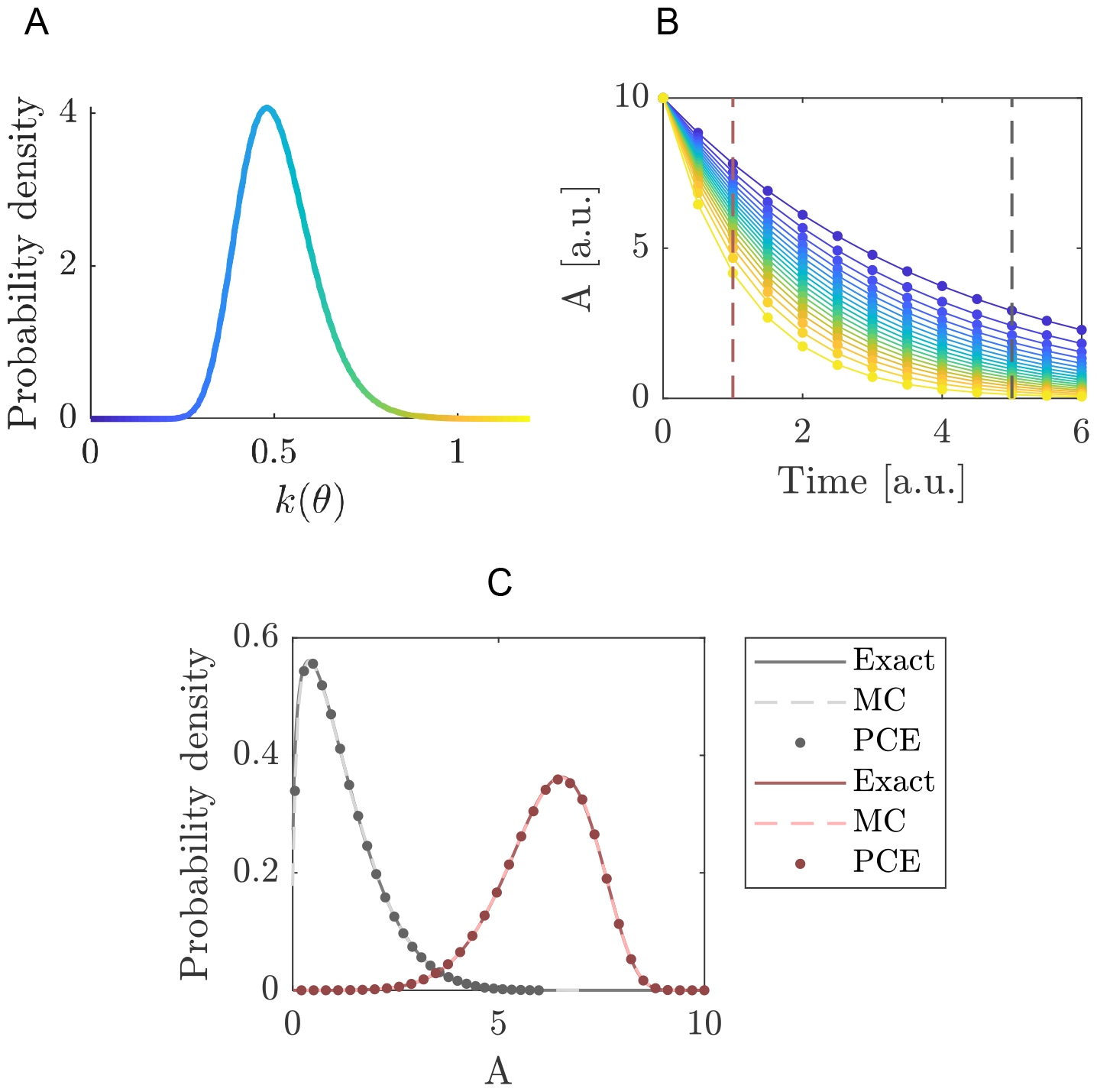
Quantifying the uncertainty propagated by the decay rate in the exponential decay model. A: Probability density function of the decay rate *k*(*θ*) with *µ* = 0.5 and *σ* = 0.2. The colour gradient corresponds to the value of *k*(*θ*). B: The concentration of *A* up to *t* = 6 seconds. The solid lines indicate the analytical solution and the dots indicate the reconstruction using PCE with Hermite basis functions and an expansion order *N* = 5. The colour for each of the solutions correspond to the colour of the line in A, which indicates the value of *k*(*θ*) used for each of the depicted solutions. C: The dashed lines in B indicate a cross-section along the model response space at *t* = 1 for the red line, and at *t* = 5 for the grey line, determined through three methods. First, the exact dynamics of the model (solid lines), second through MC sampling using the exact model dynamics (dashed lines) and finally, through MC sampling of the reconstructed function as obtained through PCE (dots).

Next, we determine how the uncertainty in *k* propagates through the model and affects concentration *A*(*t*). To that end, we expand the function *A*(*t*) = *A*_0_ exp^*−kt*^ in terms of Hermitian polynomials. Using (9), *k* is transformed into a standard normal variable *θ*. To arrive at the meta-model *Y* ^*s*^(*θ*) we truncate the expansion to a certain order *N*, as shown in (3). Choosing *N* is not straightforward and will involve some experimentation. In Figure 1 we compare results for *N* = 5 to the analytical solution for different *k* values. This shows that for this expansion order the reconstructed function accurately matches the analytical solution.

In Figure 1, we focus on the distributions of *A*(*t* = 1) and *A*(*t* = 5). This is achieved by sampling the PCE using a large sample set *χ* of reduced (i.e. standard normally distributed) variables *θ, χ*_*sim*_ = {*θ*_*j*_, *j* = 1, …, *n*_*sim*_}. The trun-cated series is then evaluated onto this sample: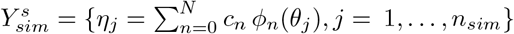 . These PDFs are obtained by kernel smoothing [44] using a sample set with *n*_*sim*_ = 10^6^, drawn from the standard normal distribution with *µ* = 0 and *σ* = 1. The kernel density estimator is given by

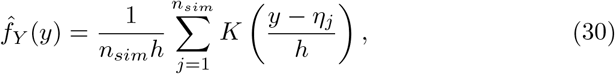

with kernel function 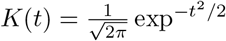 and bandwidth *h*, which is determined by Silverman’s rule of thumb [45]. Figure 1 shows that both MC and PCE perform well in reproducing the exact PDFs.

### 3.2 Example II. Biochemical reaction network: dealing with higher dimensions

In this example we present a simple model with multiple uncertainty parameters. The model describes the dynamics of two proteins *x*_1_ and *x*_2_ which bind together to form a dimer *x*_3_. We consider the following reactions:

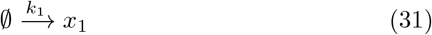

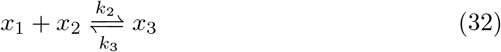

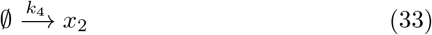

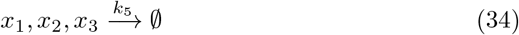

In this network, the proteins *x*_1_ and *x*_2_ are produced at rates *k*_1_ and *k*_4_. Proteins *x*_1_ and *x*_2_ reversibly bind to form species *x*_3_, with binding rate *k*_2_ and unbinding rate *k*_3_. All three proteins are degraded at the rate *k*_5_.These interactions are visualized in a reaction scheme in Figure 2. The ODEs for this system are

**Figure 2.**
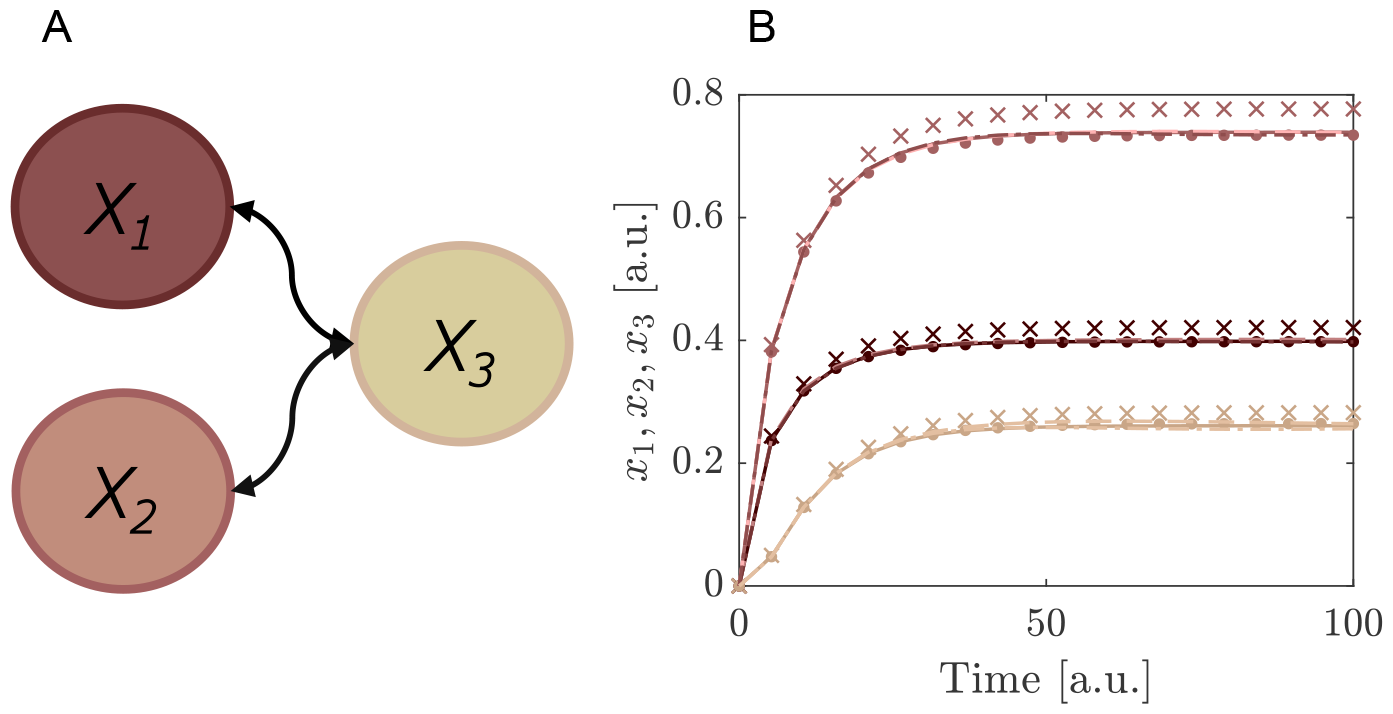
Comparing the segmented expansion with the standard non-segmented expansion. A: Interaction scheme of the model in Eqs.(35)-(37). Note that *x*_1_, *x*_2_ and *x*_3_ all have the same degradation rate and *x*_1_, *x*_2_ have production rates in the model but this is not indicated in the scheme. B: Reconstruction of the system by a Hermitian expansion. For the segmented reconstruction we used *N* = 2, *M* = 1 (crosses) and *N* = 3, *M* = 1 (dots). For non-segmented expansion the expansion order was *N* = 8 (dashed lines) and *N* = 9, (dash-dotted lines). Note that these lines overlap with the true model solution (solid lines). We used two log-normal distributions with mean and standard deviation *µ*_1_ = 0.1, *σ*_1_ = 0.1 for *k*_1_, *k*_4_, *k*_5_ and *µ*_2_ = 0.4, *σ*_2_ = 0.1 for *k*_2_, *k*_3_.

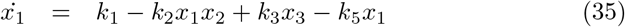

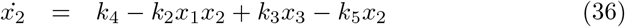

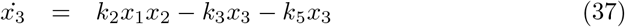

We use for the parameters *k*_1_ − *k*_5_ log-normal distributions and expand the functions *x*_1_ −*x*_3_ in terms of Hermitian polynomials. In Figure 2 we compare the results of the segmented expansion with the non-segmented expansion and the exact results. For this comparison we chose the degree of expansion and segmentation granularity such that the same number of model evaluations were required. We found that the subsequent summation to reconstruct the solutions for the differential equations improved by factors of 1000-30,000 when using the segmented expansion. See Table 1. As mentioned in the Methods section, this improvements stems from the large reduction in the number of terms to be summed over in the segmented case compared to the non-segmented expansion.

**Table 1:** Benchmarking of segmented and non-segmented expansion. An overview of the number of model evaluations *N*_*λ*_, the number of summation terms *N*_Σ_ and the time in seconds spent on summation (*t*_Σ_), for different orders *N* of expansion, both segmented *M* = 1 and non-segmented *M* = 0. The last column highlights the speed-up factor when segmentation is used (keeping *N*_*λ*_ constant).

| N | M | $N_\lambda$ | $N_\Sigma$ | $t_\Sigma[s]$ | Fold-change |
| --- | --- | --- | --- | --- | --- |
| 2 | 1 | 7776 | 3.07E+03 | 0.03 |  |
| 3 | 1 | 59049 | 1.77E+05 | 0.17 |  |
| 6 | 0 | 7776 | 1.81E+08 | 31.96 | 1065 |
| 9 | 0 | 59049 | 1.05E+10 | 5402 | 31776 |

Dividing the parameter intervals into smaller sub-intervals is a relatively straightforward and simple way to circumvent huge computation times. Other, more intricate methods have been developed to tackle models with an even larger amount of parameters [33, 46, 47]. For example, using an adaptive algorithm that is based on classical statistical learning tools can result in a “sparse” PCE, that consists of only the significant coefficients in the expansion, thereby reducing the computational cost. This method has been tested on models of stochastic finite element analysis with up to 21 parameters [46].

### 3.3 Example III. Glycolytic oscillator: mixed input PDFs and global sensitivity

Living cells obtain energy by breaking down sugar in the biochemical process called glycolysis. In yeast cells, this glycolysis was observed to behave in an oscillatory fashion, where the concentration of various intermediates were increasing and decreasing within a period of several minutes [48]. This glycolytic oscillator can be modelled as a two-component system with a negative feedback [41]:

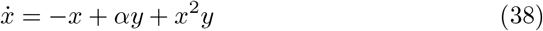

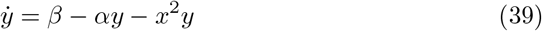

where *x* and *y* are the concentrations of ADP (adenosine diphosphate) and F6P (fructose-6-phosphate) and *α, β* are kinetic parameters. Depending on the values of *α* and *β* the system will be in a stable limit cycle or a stable fixed point [41]. In this example, we assume *α* to be uniformly distributed on the interval [0.1, 0.5] and *β* to follow a lognormal distribution with *µ* = 0.3 and *σ* = 0.1. Because both uncertainty parameters come from a different distribution, the expansion will consist of multivariate polynomials Ψ_*N,M*_ which are tensor products of the univariate polynomials. In this case, Legendre polynomials are used to expand *α*(*θ*_1_) and Hermite polynomials for *β*(*θ*_2_), where *θ*_1_ ∼ 𝒰 (−1, 1) and *θ*_2_∼ 𝒩 (0, 1). This results in a mixed polynomial for the overall expansion as exemplified with a 3rd order Legendre polynomial 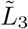 and a 3rd order Hermite polynomial 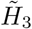, giving 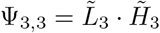 (Figure 3).

**Figure 3.**
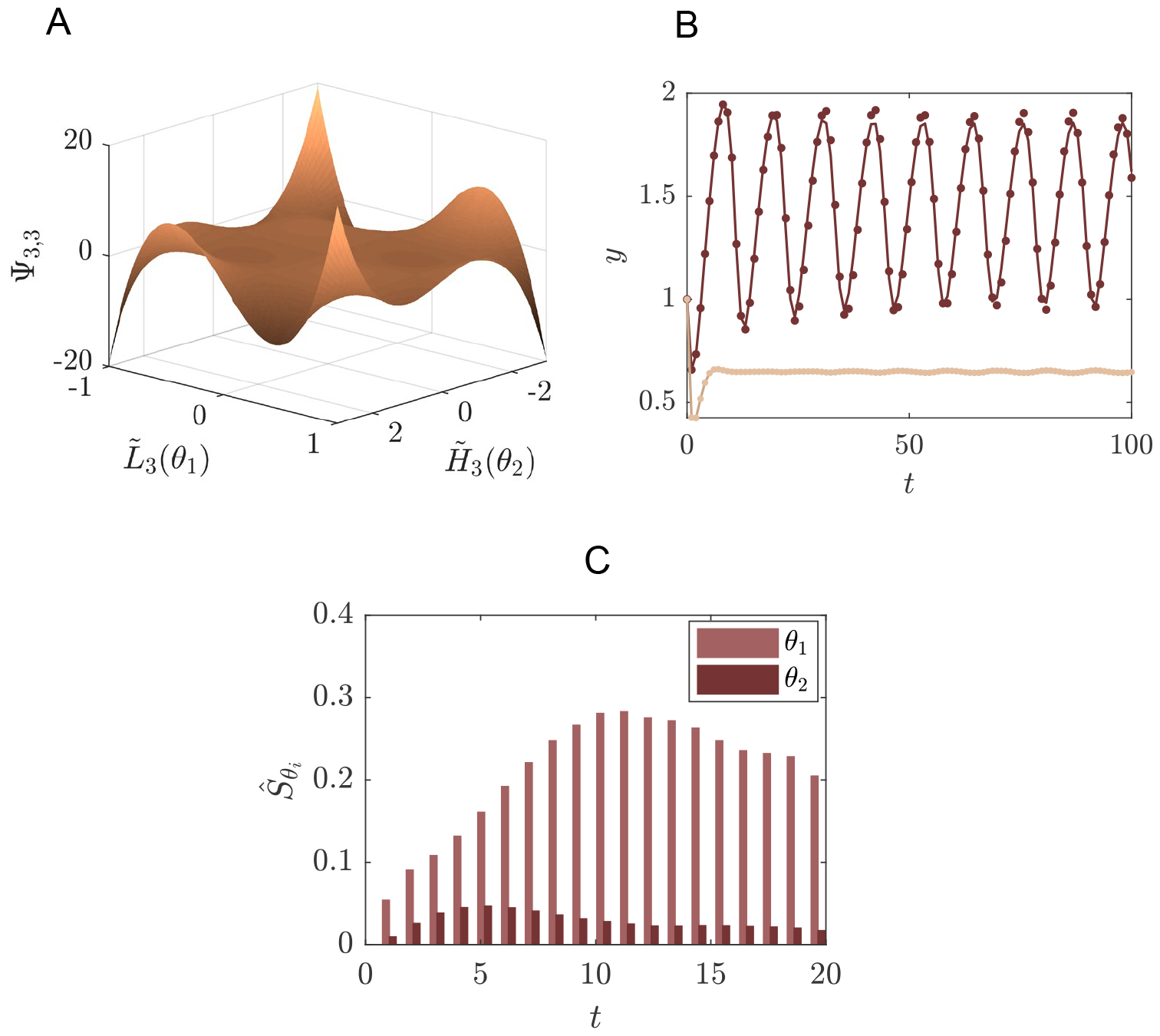
Example of a system of glycolysis as defined in (38)-(39) where PCE is performed for the concentration of species *y* and for two uncertainty parameters. A: Example of a multivariate polynomial, consisting of the tensor product between 3rd Legendre polynomial of the first random variable 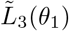 and the 3rd Hermite polynomial of the second random variable 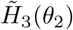. B: Solutions of concentration of fructose-6-phosphate (*y*) in the glycolytic oscillator model for two different points in the parameter space, obtained by solving the ODEs (lines) and reconstructing via PCE (dots). *α* = 0.1, *β* = 0.46 produces an oscillation (dark coloured) whereas *α* = 0.5, *β* = 0.46 gives a stable fixed point (light coloured). We used *N* = 10 as expansion order for the Legendre and Hermite polynomials. C: The first order Sobol sensitivity index 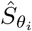 for the two random variables in the glycolytic oscillator model for the first 20 time points.

The distributions of the uncertainty parameters were chosen such that they include the bifurcation point from stable limit cycle to the stable fixed point (Figure 3). For the purpose of this example we are interested in the concentration of *y* only and therefore reconstruct this model response using PCE. A good approximation is obtained with a truncation degree of the PCE of *N* = 10.

This value is relatively high, due to the bifurcation in the system. However, this case shows that convergence can be reached using PCE despite such challenges. Yet, the computational costs are still very tractable. In the following examples we deal with cases in which still higher order expansions are necessary due to non-smooth bifurcations.

In post-processing we may use the PC coefficients to determine the first order Sobol indices for the parameters *α* and *β* at each time point, providing a representation of the global sensitivity based on variance decomposition. The Sobol indices are readily available from the PC coefficients (see Methods). They have the advantage of being global measures of sensitivity. In Figure 3 we show the first order Sobol indices given in (26) for the first and second random variable. They indicate the contribution to the total output variance of either *θ*_1_ or *θ*_2_ individually. Higher-order terms would give an indication of interaction effects between *θ*_1_ and *θ*_2_, which are also readily available from the PCE coefficients but are not considered here for brevity.

### 3.4 Example IV. Schnakenberg model: dealing with spatial discontinuities

In this example we introduce a spatial component. We consider the Schnackenberg model, which is one of the simplest, but yet realistic two-species system that can produce periodic solutions and therefore has become a prototype for reaction diffusion systems. The Schnakenberg model consists of the following (dimensionless) equations [42]

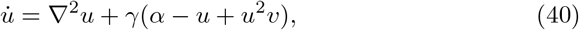

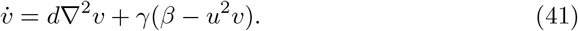

*α, β* are reaction rates, *γ* a scale parameter and *d* the ratio of diffusion constants between the species *u* and *v*. The species *u* is auto-catalytically produced by the *u*^2^*v* term in (40), whereby species *v* is consumed. There are certain combinations of the parameters *α, β, γ* and *d* for which the system will exhibit a stable pattern [49]; this region of parameter space is called the Turing space (TS). For the purpose of this example we limit the number of uncertainty parameters to one: the parameter *α*, fixing the other parameters at *β* = 1, *γ* = 5 and *d* = 20. We assume *α* to be distributed as *α*∼𝒰 (0.001, 0.45) and determine the TS for a range of *α* (Figure 4) using linear stability analysis (for details see [42, 49]). To that end, the model is simulated on a 1D grid of 20 cells. We focus on the concentration of species *v* at steady state and consider an expansion by both Legendre polynomials and Haar wavelets.

**Figure 4.**
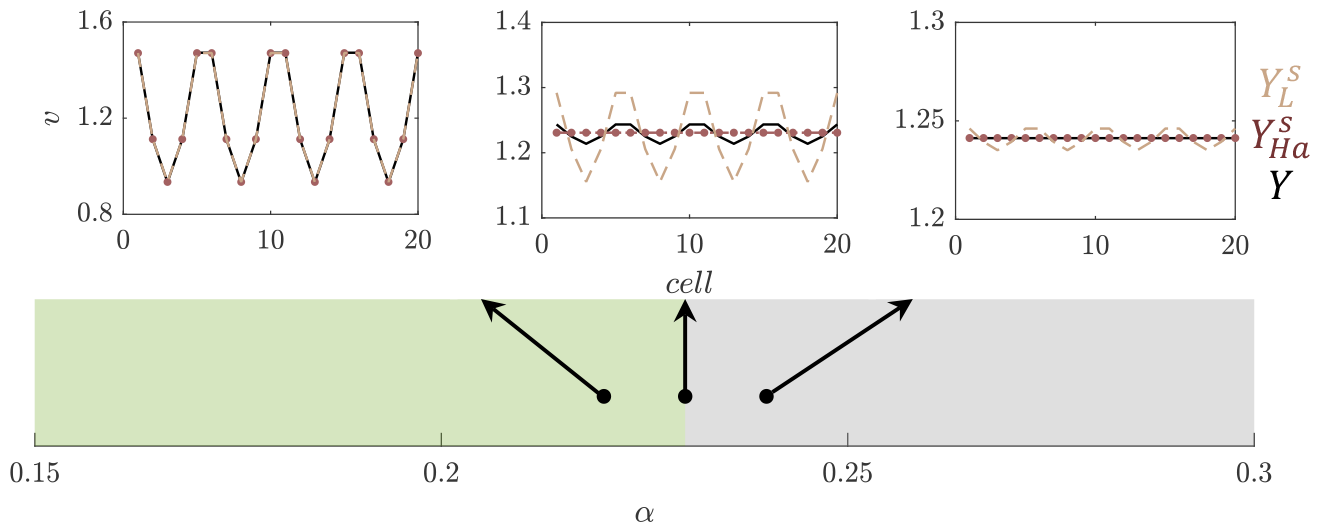
Reconstruction of the concentration of species *v* of the Schnakenberg model, as defined in (41) at steady state, comparing Legendre and Haar wavelet expansions. We consider the patterning of *v* in the Schnakenberg model for *α*∈[0.15, 0.3], as indicated on the main x-axis and assuming *α*∼𝒰 (0.001, 0.45), while fixing *β* = 1, *γ* = 5 and *d* = 20. The colour along this axis indicates the region in the 1D-parameter space whether a stable pattern will form (i.e., *α* is inside the Turing Space (TS), indicated by green colouring), or a homogeneous spatial distribution of *v* (grey colour). We highlight three examples of the patterns formed for different values of *α*, one inside the TS (left inset, *α* = 0.22), one close to the boundary (middle inset, *α* = 0.23) and one outside the TS (right inset, *α* = 0.24). Within these examples we compare the true solution *Y* (solid black line) to the reconstructed function of *v* by PCE in terms of Legendre polynomials 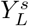 by a segmented expansion with polynomial order *N* = 18 and segmentation granularity *M* = 3 (dashed line), or Haar wavelets 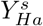 (dots), using the first 128 wavelets in the expansion,i.e. *N* = 6 resolution levels.

Polynomial chaos expansion is known for being inaccurate in regions that contain discontinuities [8, 31, 32]. In this example, the lack of convergence in PCE can be seen along the boundary of the patterning space (TS) in Figure 4, where the expansion by Legendre polynomials is indicated with the dashed lines.

For the reconstruction of concentration of *v* in terms of Legendre polynomials, we used a segmented expansion with *N* = 18 and a segmentation granularity of *M* = 3, leading to a total of 126 model evaluations used in the expansion. To show that Haar wavelets perform much better in such a region, we additionally do an expansion in terms of Haar wavelets. As resolution level we take *N* = 6, which means a total of *N*_*w*_ = 128 wavelets are used in the expansion. In Figure 4 the performances of Legendre polynomials and Haar wavelets are compared in the vicinity of *α* = 0.23 (middle inset, Figure 4), showing that the Haar-wavelets provide an improvement in accuracy at the bifurcation point, while using the same number of model evaluations (i.e. the same amount of information and computational cost) for the expansion.

### 3.5 Example V. Trichome patterning: dealing with spatial discontinuities

As an extra example of pattern formation we consider a model that describes trichomes. Trichomes are hairs found on the epidermal layer of leaves. In *Arabidopsis Thaliana* these trichomes form a regular pattern, where each trichome is separated by around three to four epidermal cells [50]. The model studied here consists of three proteins and their interactions which, taken together, can explain features of trichome patterning [43, 51]. Protein transpararant testa glabra1 (TTG1) binds to the transcription factor glabra3 (GL3) which together form a trichome-promoting complex, called the activating complex (AC) [43]. Experimental data suggests that TTG1 is depleted from cells neighbouring a trichome [43]. For this reason the interaction between TTG1 and GL3 is modelled in a substrate-depletion form (Figure 5), where TTG1 acts as a substrate for the formation of AC [43]. After non-dimensionalisation this model consists of four parameters, none of which have been experimentally determined, highlighting the substantial amount of uncertainty within this model [52, 53]. Here, we examine the propagation of uncertainty in the parameters to the predicted pattern.

**Figure 5.**
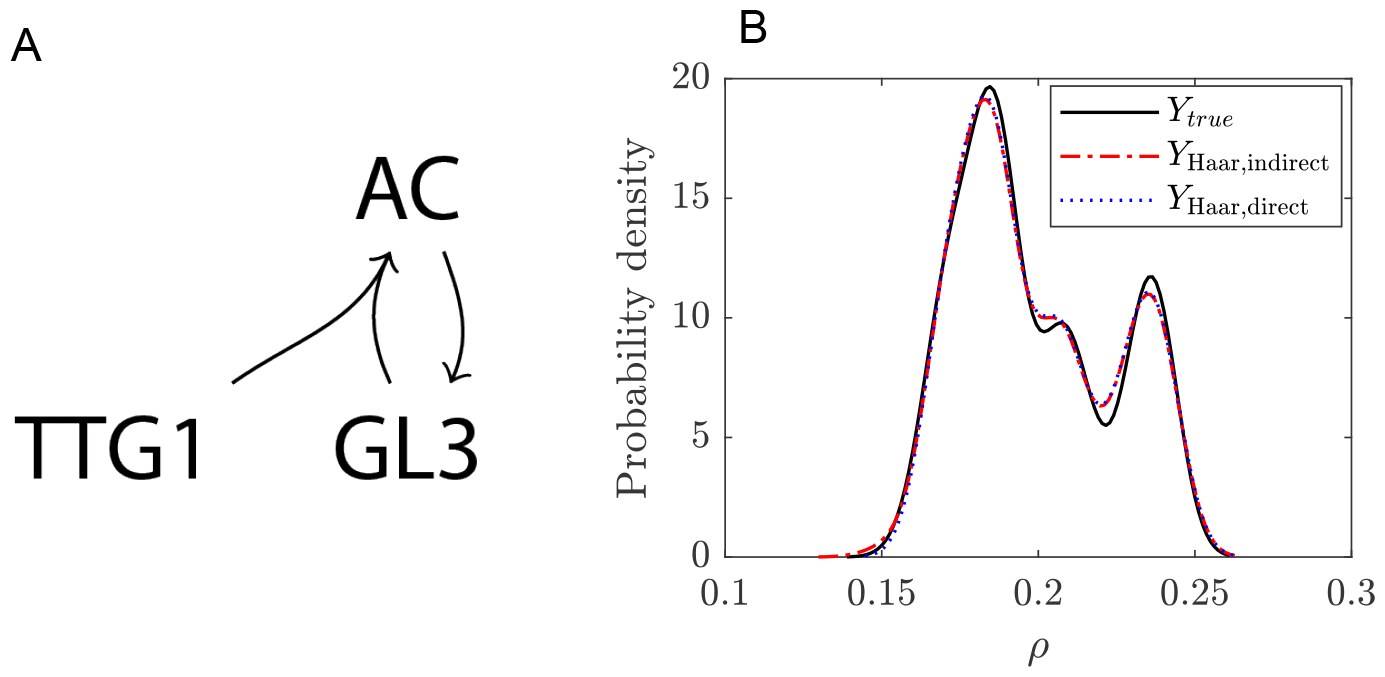
Uncertainty quantification for the trichome system. A: Schematic of the model. B: Probability density function of the trichome density in the Turing Space using either the indirect (dashed red line) or direct expansion (dotted blue line) method and for comparison the solution of the real model (solid black line). A resolution level of 3 (i.e., a total of 16 wavelets) has been used for the expansion.

The trichome patterning is described by the following set of coupled ODEs

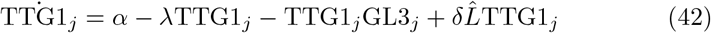

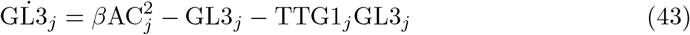

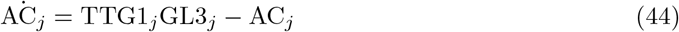

where *α, λ, δ* and *β* are parameters in the model and 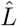 describes the coupling between the cells. The subscript *j* indicates the *j*^th^ cell. We solve these equations for 400 cells, grouped on a hexagonal grid of 20 by 20 cells.

In this example we focus on the parameter *α*, the basal production for TTG1. We assume this parameter to be uniformly distributed on the interval [0.4, 0.9]. We are interested in the number of trichomes that are predicted by the model, therefore we consider the trichome density *ρ* (total number of trichomes divided by the total number of cells in the simulated tissue) as the model response of interest. The number of trichomes is determined by simulating the system until steady state is reached and counting the number of cells for which the concentration of AC exceeds a threshold. The amount of AC is considered to be an indicator for trichome cell fate in plants, however the biological threshold for this is unknown. We set this threshold to the half-maximum of AC in the system. This leads to the following description of trichome density:

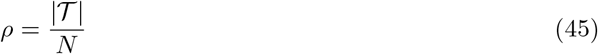

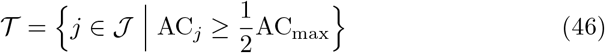

where 𝒯 is the set of cells which exceed the AC threshold, 𝒥 is the set of all cells on the grid, *N* is the total number of cells and |𝒯 | is the cardinality of 𝒯 .

The goal is to determine the uncertainty in *ρ* as a result of the uncertainty in *α*. To this end, we employ two different approaches. For both approaches we first transform *α* to a standard uniform variable by *α* = *T*^*−*1^(*θ*), using the transform function for a uniform variable given in (8). The first approach, refered to as the indirect approach, is the same as used in Example IV. To reconstruct the concentration at steady state for all cells, we expand the concentration of AC using Haar wavelets. From the result we may determine *ρ*. In this process we discriminate between cases where there is a pattern and where there is no pattern. For the latter, we need not solve the system as *ρ* = 0. Through linear stability analysis we determine beforehand whether a pattern will form or not, i.e., whether the chosen parameter set is in the Turing Space (TS) [42]. For a certain realisation *θ* we can determine *ρ* by

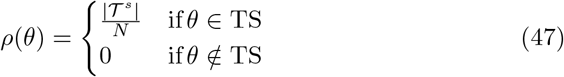

where 𝒯^*s*^ is the set of trichomes as determined from the reconstructed AC concentration profile.

Our second, direct approach is to directly reconstruct *ρ* as

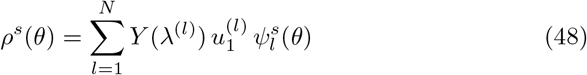

Similarly, as we did for *ρ*(*θ*) we can define *Y* (*λ*^(*l*)^) as

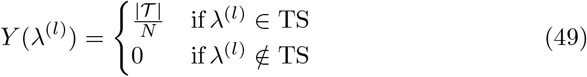

In other words, we only solve the system and determine the trichome density if the parameter set falls within the Turing space. This lends robustness to the PCE for the non-smooth parts of the function *Y* (*θ*) and at the same time limits the amount of simulations to be performed, as the non-patterning parameter combinations need not be solved for.

We show here that there are multiple ways in which the uncertainty in the output can be captured. In this case of trichome patterning, we first tested an indirect method where the model output consists of concentration profiles from which the pattern features have to be extracted in post-processing, and secondly, we showed that the pattern feature could also be expanded directly, by taking the density as model output. Comparing the indirect and direct approaches, we conclude that both have similar levels of accuracy (Figure 5). Note that in both cases the expansions converge to the real solution at resolution level N = 6, which means a summation of 128 wavelets. The PDF in Figure 5 is constructed using 10^3^ samples which costs 4.7 seconds for the wavelet reconstruction as opposed to 80.9 seconds for solving the full model a 1000 times in an MC approach.

### 3.6 Example VI. Plasmid transfection: dealing with correlated parameters

It can happen that in parameter space a structure occurs, i.e., that the multivariate joined probability cannot be written as a product of univariate distributions. In this section we exemplify how to handle such a case. As an example we choose the transfection of mammalian cells (e.g., Human embryonic kidney cells, HEK293) with two plasmids: plasmid *pl*_1_ with the construct for induction (e.g. through chemical or optogenetics [54]), and plasmid *pl*_2_ with the reporter construct. Denoting by *n* the number of *pl*_1_ plasmids and by *m* the number of *pl*_2_ plasmids in a specific cell, the corresponding reaction scheme reads:

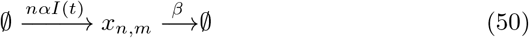

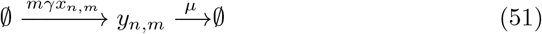

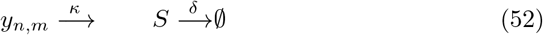

where *x*_*n,m*_ and *y*_*n,m*_ are the concentrations of molecule x and y, resp., in a bacteria with plasmid composition *nm. S* is the concentration of the reporter molecule in the bulk. *α* and *γ* are the corresponding production rates per plasmid and *β, µ* are the degradation rates. *I*(*t*) is the external induction signal. In what follows we will set *I*(*t*) = 1 for *t*≥0.

For sake of clarity we keep the system simple and ignore all complicating effects, such as gene expression noise, maturation of the reporter construct (e.g. GFP). However, for transient transfections of the mammalian cells, we need to take into account the distribution of plasmids among the cells, which is not constant due to cell division. The set of Ordinary Differential Equations describing the reactions given by 50-52 read:

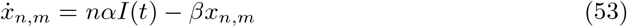

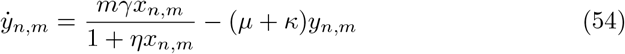

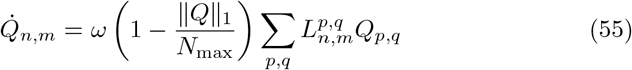

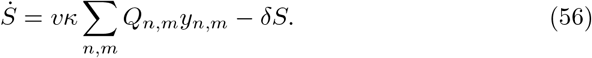

The parameter *η* captures the saturation of the expression capacity of the cells. We set *η* = 0.2 *V* ^*−*1^ throughout the calculations. *V* is the unit of the volume of the cells, e.g., *V* = *µm*^3^. *Q*_*n,m*_ is the number of cells that have *n, m* plasmids which are growing by *ω*. To follow the partition of plasmids upon division, the tensor *L* is introduced and described in further detail below.∥*Q*∥_1_ = Σ*n,m Q*_*n,m*_ = *N*_tot_ which is the total number of cells at time *t* and *N*_max_ is the maximum number of bacteria due to environmental constraints. The growth rate is set to *ω* = 0.034 *h*^*−*1^ based on a doubling time of 20 hours as being measured for HEK293T cells [55, 56]. *S* is the reporter (e.g. SEAP [57]) accumulated in the bulk due to secretion by the cells with rate *κ*. For non-secreted reporters (e.g. YFP) for single cell measurements, *κ* is set to zero. *v* is a volume correction factor, set to *v* = 10^*−*8^.

We assume that the rate of dilution of the plasmids corresponds to the growth rate of the cells, i.e. plasmids are lost upon division. In reality, the plasmids are also degraded, but we assume the time scale of this degradation is much longer than the experimental duration and can therefore be ignored. To model the partition of plasmids upon cell division we assume that any partition of the number of plasmids is possible. The probability of finding a certain partition is given by the binomial distribution

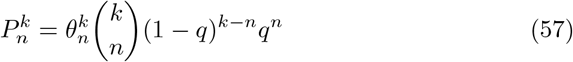

where *n* is the number of plasmids in the mother cell, *k* the number of plasmids in one of the daughter cells and *q* the probability, which for equal cell division is *q* = 0.5. 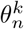 is the discrete unit step function with 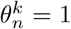 for *k* ≥ *n* and 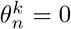 else. We can write the rate of change in the pools of cells as

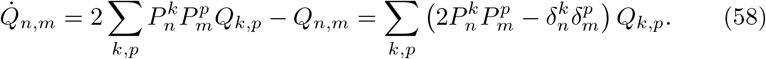

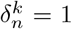 for *k* = *n* and 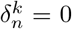 else. From this, we can define the tensor used in Eq. 55:

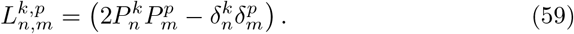

We treat the plasmid uptake as a Poisson process, i.e., the number of plasmids inside a bacteria is Poisson distributed. Assuming that the mean number of plasmid taken up is the same for *pl*_1_ and *pl*_2_ and a correlation exists between the uptake of two plasmids, the distribution is given by [58]:

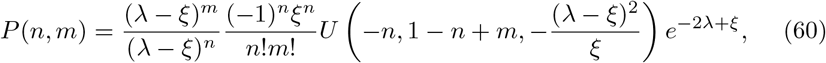

where *U* is Kummer’s confluent hypergeometric function, *λ* is the mean number of plasmids taken up, and 0 ≤ *ξ* ≤ *λ* is the correlation parameter. Note that for *ξ* → 0 we find:

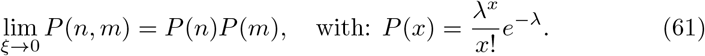

The correlation between *n* and *m* reads:

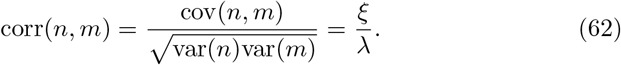

In Fig. 6A we show *P* (*n, m*) for *λ* = 3 and *ξ* = 2.4. The asymmetry due to the correlation corr(*n, m*) = 0.8 can be clearly seen. The orthonormal basis function *ϕ*_*p*_(*n*) with respect to the Poisson distribution read [59]:

**Figure 6.**
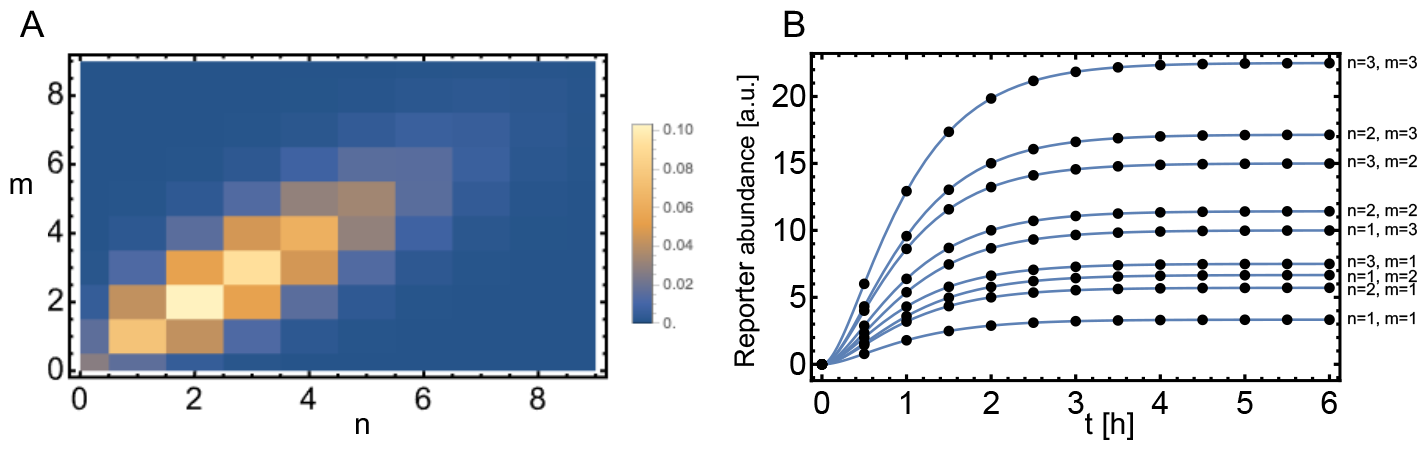
Expansion with respect to a bivariate Poisson distribution. A: Bivariate Poisson distribution given by Eq. 60 with *λ* = 3 and *ξ* = 2.4. B: Comparison of the exact solution for *y*_*n,m*_(*t*) given by Eq. 54 (solid lines) to the spectral expansion with *N* = 6 (dots). Parameters: *α* = 2 (*V h*)^*−*1^, *β* = 2 *h*^*−*1^, *γ* = 6 *h*^*−*1^, *η* = 0.5 *V* ^*−*1^, *µ* = 1.5 *h*^*−*1^, *κ* = 0 *h*^*−*1^.

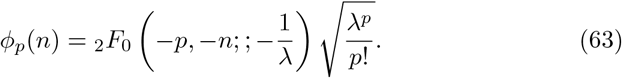

The matrix 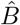 defined in Eq. 4 can be calculated analytically and is given by:

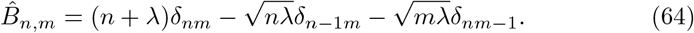

Note the dependence on the parameter *λ*.

We first consider a system in which the mammalian cells do not secrete the reporter molecule, e.g., YFP. In this case we set *κ* = 0, and consequently *S*(*t*) = 0, ∀*t*. In Fig. 6B we show *y*_*n,m*_(*t*) for different plasmid compositions and compare the expansion of *y*_*n,m*_(*t*) of the order *N* = 6 (dots) to the exact results (solid lines). A question of interest is what is the measured distribution of fluorescence intensities of the mammalian cells. The distribution of reporter molecule abundance is given by:

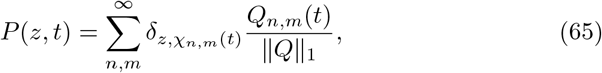

where *χ*_*n,m*_(*t*) is a binning function: *χ*_*n,m*_(*t*) = *g* ⌊*y*_*n,m*_ (*t*)*/g*⌋ (other binning methods are also possible, of course). Note that for sake of simplicity we ignore the transient phase after cell division in which the dynamic of *y*_*n,m*_ adopts to the new, reduced plasmid composition. This is also a reasonable simplification considering that the steady state of *y*_*n,m*_ is roughly reached after 4 *h*, as can be seen in Fig. 6B, in contrast to a cell doubling time of 20 h. In Fig. 7A we show the distribution *P* (*z, t* = 50) for *g* = 5 for *ξ* = 2.4 (left) and *ξ* = 0 (right); we used the expansion with *N* = 6 for the calculations in Eq. 65.

**Figure 7.**
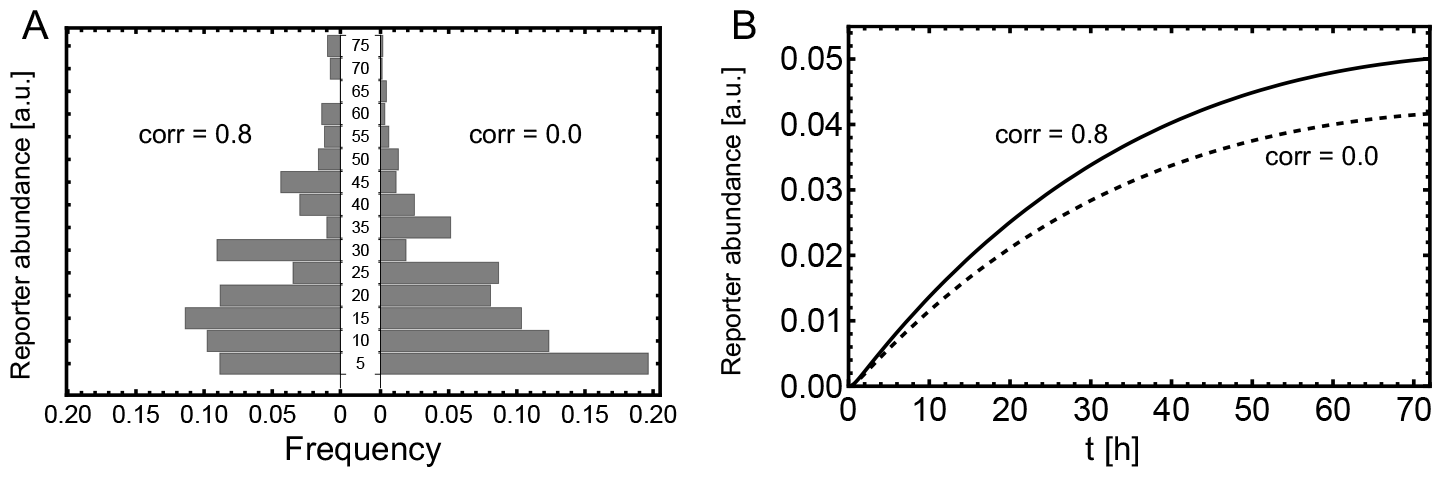
Measurement of the reporter. Comparison of correlated and un-correlated plasmid uptake. A: Histogram of the single cell readout, measured 6 h after induction. The parameters are the same as stated in Fig. 6 and the histogram is calculated according Eq. 65 with *g* = 5 B: Reporter abundance in the bulk measured for three days. The parameters are the same as in A, besides *κ* = 3 *h*^*−*1^, *δ* = 0.01 *h*^*−*1^ and *v* = 10^*−*8^. We used for the calculations the expansion of *y*_*n,m*_ with *N* = 6.

Next, we consider a segregated reporter, e.g., SEAP. In this case the reporter abundance in the bulk is measured instead of single cell measurements. The temporal evolution of the bulk reporter is governed by Eq. 56. In Fig. 7B one can see that the dynamics of the reporter is clearly different due to correlation of the plasmid uptake.

### 3.7 Benchmarking of MC and PCE

To show the advantage of using PCE over MC, we turn to a trichome model that consists of 8 states and 31 parameters. We simulate this model on a grid of 35-by-35 hexagonal cells, resulting in a system of 9800 coupled ODEs [43]. We fix the initial conditions to the solution of the homogeneous steady state of a ‘default’ parameter set. We also use this parameter set to set the distributions for the uncertainty parameters, by choosing the mean to be the default value and the standard deviation to be 5% of the mean. In this benchmarking example, we aim to determine the mean and variance of the trichome density and do so by Monte Carlo simulation (as described in Methods) and PCE in terms of Hermite polynomials, with a varying number of uncertainty parameters. We start by assuming the first parameter *θ*_1_ to following a lognormal distribution (see appendix for parameters and model equations), then in a second comparison we additionally assume *θ*_2_ to follow a lognormal distribution (with different mean and variance), up to *θ*_1_, …, *θ*_5_. For each of the different number of uncertainty parameters we determined the mean and variance, and compared the time computational time between MC and PCE (Figure 8). For both PCE and MC we used parallelisation, by running the model simulations on 9 cores in parallel (3.1 GHz CPU). We found convergence for MC at 1000 samples, which on average took between 75 to 80 minutes. For PCE we find a variety of computational times, ranging from around 2 minutes to 56 minutes for the lowest and highest number of uncertainty parameters, respectively. Note that for PCE a major computational advantage is the low cost of calculating the mean and variance. The mean follows directly from the zeroth coefficient: ⟨*Y* ⟩ = *c*_0_ and the variance is given by:

**Figure 8.**
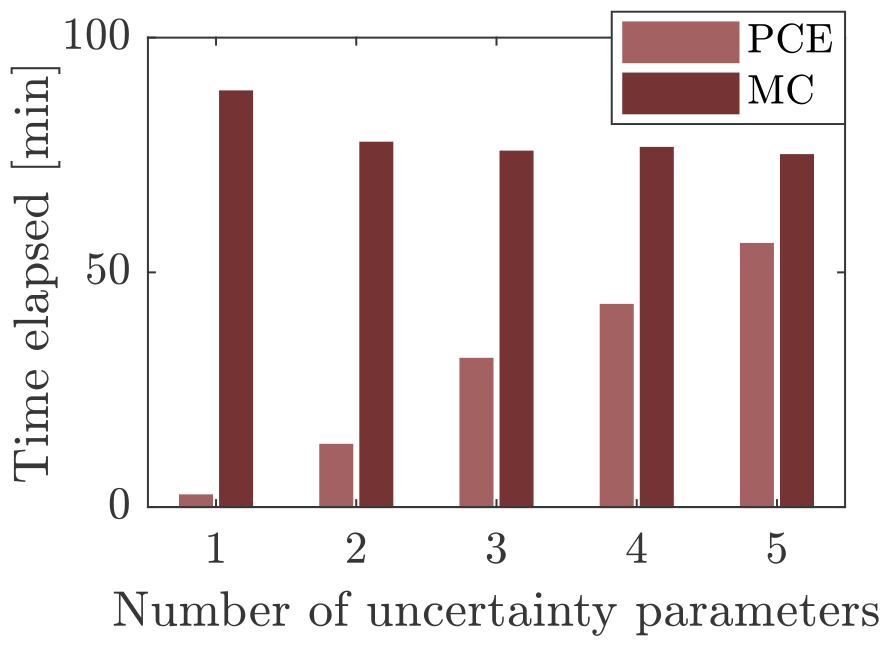
Benchmarking of PCE and MC. An overview of the computational times spent for PCE (light red) and MC (dark red).

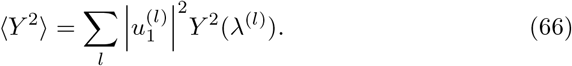

In comparing the mean and variance of the trichome density we find good agreement between MC and PCE (Table 2).

**Table 2:** Comparison of mean and variance found by MC and PCE. NPar indicates the number of uncertainty parameters.

| NPar | PCE order | $\langle Y \rangle$ (MC) | $\langle Y \rangle$ (PCE) | $\langle Y^2 \rangle$ (MC) | $\langle Y^2 \rangle$ (PCE) |
| --- | --- | --- | --- | --- | --- |
| 1 | 30 | 0.033 | 0.033 | 3.80E-04 | 3.90E-04 |
| 2 | 10 | 0.034 | 0.031 | 6.15E-04 | 5.44E-04 |
| 3 | 7 | 0.034 | 0.030 | 7.26E-04 | 6.36E-04 |
| 4 | 5 | 0.034 | 0.030 | 7.19E-04 | 6.18E-04 |
| 5 | 4 | 0.034 | 0.029 | 7.22E-04 | 6.29E-04 |

## 4 Discussion and Conclusions

Through a series of examples we have presented an efficient and widely-applicable version of spectral methods for quantification of the effect of parameter uncertainty on model outcomes. The present scheme utilizes non-intrusive spectral projection based on polynomial functions or wavelets. The orthonormal properties of those functions provide a novel scheme to determine the expansion coefficients in a computationally fast way. The scheme is similar to the Golub-Welsh algorithm known from Gaussian quadrature [60]. In fact, the points indicating the roots of the polynomials used obtained in quadrature procedures, exactly correspond to the eigenvalues obtained from the matrix, which plays a central role in our approach. Quadrature methods as well as sampling methods are traditionally used to determine the coefficients of expansions of functions one is interested in [6]. The approach used here does effectively the same and provides an alternative to existing approaches, with the advantage that it is flexible and applicable to any set of orthonormal basis functions.

The method presented here requires no modification of the model equations. This is in general the main advantage of non-intrusive methods: there is no need to recast the model into a probabilistic framework. Instead, the random behaviour of parameters is accounted for through a set of deterministic simulations of the process for a restricted number of parameter values. These values are chosen such that they reflect the uncertainties in the parameters. To test the performance of our method we applied it to six different models: (1) a model of exponential decay, (2) a biochemical reaction network, (3) the glycolytic oscillator, (4) the Schnakenberg model, (5) a trichome model, and (6) transient mammalian plasmid uptake. Model (4) and (5) deal with spatial pattern formation. For each test case, the results of the PCE were compared to, if available, analytical solutions, non-PCE numerical simulations, or Monte Carlo simulations. In these comparisons we mostly focused on the accuracy of PCE. Even though the computational advantage is one of the major reasons for using PCE techniques, we do not focus on it because this aspect has already been extensively explored in various applications. For such comparisons we refer to other publications, see for example [6, 61, 62]. The logic behind the compututational advantages of PCE over MC extends to the methods we have presented here and is evident, e.g., in the case of trichome pattern formation (Example V), for which the output of the model was obtained 17 times faster when using PCE instead of MC.

The accuracy of the reconstruction by PCE depends on the choice of expansion order and basis functions. While the latter is determined by the PDFs of the input parameters, the choice of expansion order has to be chosen by the user. For example, in Examples I and III we chose *N* = 5 and *N* = 10, respectively. These choices were based on careful observation of the convergence properties of the method. In some cases the expansion order has to be chosen prohibitively large. For such a situation we propose an alternative approach that segments the parameter interval into subintervals, essentially zooming in on these sub-intervals such that a lower expansion order can be used in each sub-interval. In Example II we have shown that this segmentation approach can greatly reduce the computational costs in summation part of the expansion, thus providing a way to circumvent the curse of dimensionality. Such adaptations are required for the more difficult high-dimensional cases and the segmentation is a relatively simple and straight-forward method to tackle dimensionality problems yielding a piecewise continuous approximation to the original function. It is an alternative for so-called sparse PCE methods, that utilize only a small subset of the polynomial basis functions in order to limit the amount of model evaluations [33, 46, 47].

Convergence of the PCE may be poor in regions of the parameter space around a bifurcation [7]. In PCE, smooth polynomials are used in the expansions and they may show effects similar to the Gibbs phenomenon in Fourier expansions, i.e., the spectral basis is not suitable and leads to a slow convergence.

Since smooth functions like the Hermite and Legendre polynomials will fail to describe steep or discontinuous solutions, we explored the use of Haar wavelets. These wavelets lead to localized decompositions and this produces more robust behaviour [32]. We show in Example IV (Schnakenberg model) the advantages of using Haar wavelets over polynomials by focusing on the region in parameter space where the system transitions from spatially heterogeneous dynamics to spatially homogeneous dynamics take place. Around this bifurcation point an expansion in terms of Haar wavelets also shows slow convergence, but the accuracy of the expansion is 10 times better than when Legendre polynomials are used. For a fair comparison we kept the number of model evaluations in both approaches the same. In the vicinity of bifurcations Haar wavelets show greater robustness than the traditional polynomial basis functions. They thus provide a useful tool for biological systems which often feature such discontinuity.

In Example V (trichome pattern formation) we have highlighted the flexibility of the method: certain quantities, e.g., the scalar quantity of trichome density, can either be directly expanded or indirectly. By making use of that adaptability the number of model evaluations can be reduced while maintaining the same level of accuracy.

Finally, in Example VI we illustrate how to handle correlated parameters by means of correlated plasmid uptake by mammalian cells. We show how single-cell or bulk readout can be calculated using spectral expansion.

For a lower number of uncertainty parameters we find only small differences between the mean and variance determined by MC and PCE. As the dimensionality of the problem increases, PCE loses accuracy while showing increasingly higher computational times (Figure 2). Following the trend in Figure 2 for 6 uncertainty parameters, MC would likely be slightly faster. For such high-dimensional problems, the computational cost for PCE could be improved by e.g. sparse PCE [33, 46, 47], which has not been covered here for brevity but could be implemented within the presented scheme just as well. Furthermore, some accuracy could be gained by using Haar-wavelets instead of Hermite polynomials, which might improve the results around bifurcations in a patterning model such as the one presented here.

Overall, the approach presented here consists of a number of easy-to-implement steps and is widely applicable to a variety of systems which would be computationally costly when used in the context of uncertainty quantification. We believe that this approach provides a valuable tool in the tool-box for computational systems biology.

## 6 CRediT authorship contribution statement

C.F. conceived the project, A.D. performed all calculations, A.D., J.M. and C.F. wrote the manuscript.

## 8 Data availability

The code is publicly available at: https://github.com/AnnaDeneer/SpectralExpansion.

