## Supplement for "Spectral methods for prediction uncertainty quantification in Systems Biology"

### 1 Supplementary information

#### 1.1 S1: Expanding a meta model

To illustrate the derivation of (5) and (6), we consider here the case with only one uncertainty parameter $\theta$, distributed according to PDF $P(\theta)$. The involved Hilbert space consists of functions of $\theta$ that are square-integrable with respect to the inner product

$$\langle f, g \rangle = \int f(\theta)\, g(\theta)\, P(\theta)\, d\theta \tag{1}$$

for any two functions $f(\theta)$ and $g(\theta)$ in this space. So, $P(\theta)$ acts as weight function in the inner product.

In this Hilbert space we make use of a set of basis functions $\phi_n(\theta), n = 0, 1, 2, \ldots$ that are orthonormal with respect to this inner product. These may be Legendre, Hermite, Jacobi, Chebyshev, Laguerre, and other polynomials, but also Bessel functions, Hankel functions, wavelets and other functions may be used. The only requirement is orthonormality.

We assume the model response $Y(\theta)$ to be in the Hilbert space. This implies that we may expand $Y(\theta)$ in terms of the basis functions:

$$Y(\theta) = \sum_{n=0}^{\infty} c_n\, \phi_n(\theta), \tag{2}$$

with the expansion coefficients given by

$$c_n = \langle Y, \phi_n \rangle = \int Y(\theta)\, \phi_n(\theta)\, P(\theta)\, d\theta. \tag{3}$$

It is convenient to associate with $Y(\theta)$ the matrix $\hat{Y}$ whose elements read as

$$\hat{Y}_{n,m} = \langle \phi_n, Y \phi_m \rangle. \tag{4}$$

Assuming $Y$ to be an analytical function in $\theta$, we may write down its Taylor expansion:

$$Y(\theta) = \sum_{k=0}^{\infty} d_k\, \theta^k. \tag{5}$$

For the matrix $\hat{Y}$ this implies that it is given by the expansion

$$\hat{Y} = \sum_{k=0}^{\infty} d_k\, \hat{B}(k), \tag{6}$$

with the elements of matrix $\hat{B}(k)$ given by

$$\hat{B}_{n,m}(k) = \langle \phi_n, \theta^k\, \phi_m \rangle. \tag{7}$$

Using the completeness relation of the basis functions

$$\sum_{n=0}^{\infty} \phi_n(\theta)\, \phi_n(\theta')\, P(\theta) = \delta(\theta - \theta'), \tag{8}$$

we may factorize this matrix. For $k = 0$ we have $[\hat{B}(0)]_{n,m} = \delta_{n,m}$, and for $k \geq 1$

$$\hat{B}(k) = \hat{B}^k, \tag{9}$$

where $\hat{B} \equiv \hat{B}(1)$. So, $\hat{Y}$ can be written as

$$\hat{Y} = \sum_{k=0}^{\infty} d_k\, \hat{B}^k. \tag{10}$$

with

$$\hat{B}_{n,m} = \int \phi_n(\theta)\, \theta\, \phi_m(\theta)\, P(\theta)\, d\theta. \tag{11}$$

$\hat{B}$ is a symmetric, $(\infty \times \infty)$ matrix. So, it has real eigenvalues $\lambda^{(l)}, l = 1, 2, \ldots$ and there exists an orthogonal basis of $\infty$-dimensional eigenvectors $u^{(l)}$. Note that, as usual in the literature, but in contrast with the basis in the Hilbert space introduced above, the counting here starts at $l = 1$. The completeness relation for the $u^{(l)}$ reads as

$$\sum_{l=1}^{\infty} u_i^{(l)} u_j^{(l)} = \delta_{i,j}. \tag{12}$$

Using the relations obtained above, we may rewrite the expansion for $Y$ in the following way:

$$\begin{aligned}
Y(\theta) &= \sum_{n=0}^{\infty} \hat{Y}_{0,n}\, \phi_n(\theta) && (13)\\
&= \sum_{n=0}^{\infty} \sum_{k=0}^{\infty} d_k\, (\hat{B}^k)_{0,n}\, \phi_n(\theta) && (14)\\
&= \sum_{n=0}^{\infty} \sum_{k=0}^{\infty} \sum_{i=0}^{\infty} \sum_{l=1}^{\infty} d_k\, (\hat{B}^k)_{0,i}\, u_{i+1}^{(l)}\, u_{n+1}^{(l)}\, \phi_n(\theta) && (15)\\
&= \sum_{n=0}^{\infty} \sum_{k=0}^{\infty} \sum_{l=1}^{\infty} d_k\, (\lambda^{(l)})^k\, u_1^{(l)}\, u_{n+1}^{(l)}\, \phi_n(\theta) && (16)\\
&= \sum_{n=0}^{\infty} \sum_{l=1}^{\infty} Y(\lambda^{(l)})\, u_1^{(l)}\, u_{n+1}^{(l)}\, \phi_n(\theta) && (17)\\
&= \sum_{l=1}^{\infty} Y(\lambda^{(l)})\, u_1^{(l)} \psi_l(\theta), && (18)
\end{aligned}$$

In the last step we used an orthogonal basis transformation from $\phi(\theta)$ to $\psi(\theta)$ with the help of the orthogonal vectors $u^{(l)}$:

$$\psi_l(\theta) \equiv \sum_{n=0}^{\infty} u_{n+1}^{(l)} \phi_n(\theta). \tag{19}$$

The important conclusion is that we may express the model response $Y(\theta)$, which originally depends on the continuous parameter $\theta$, in terms of discrete values $Y(\lambda^{(l)}), l = 1, 2, \ldots$. This implies that $Y$ needs to be evaluated only at the points $\theta = \lambda^{(l)}, l = 1, 2, \ldots$ in parameter space. Note that most ingredients of the formalism can be calculated in advance and once and for all.

The corresponding meta-model response $Y^s$ we are aiming at is obtained from $Y$ by taking into account only the lowest $N$ basis functions, for some integer $N$. So,

$$Y^s(\theta) = \sum_{n=0}^{N} c_n \, \phi_n(\theta). \tag{20}$$

Note, that the coefficients $c_n, n \leq N$, do not change if we omit the terms with $n > N$, thanks to the orthonormality of the basis functions. For the meta-model we have

$$Y^s(\theta) = \sum_{l=1}^{N} Y(\lambda^{(l)}) \, u_1^{(l)} \, \psi_l^s(\theta). \tag{21}$$

We remark that omitting higher order basis functions has also consequences for the $\psi_l$. Instead of $\psi_l$ we now use

$$\psi_l^s(\theta) \equiv \sum_{n=0}^{N} u_{n+1}^{(l)} \, \phi_n(\theta). \tag{22}$$

In practice one should for each case analyze the accuracy of $Y^s$ in approximating $Y$, especially as a function of $N$.

#### 1.2 S2: Handling correlated parameters

Let $P(\theta_1, ..., \theta_M)$ be the distribution for $M$ correlated parameters, such that $P(\theta_1, ..., \theta_M) \neq \prod_{i=1}^{M} P_i(\theta_i)$, where $P_i$ is the marginalised distribution for $\theta_i$. We can construct functions that are orthonormal with respect to the joined distribution $P$ by:

$$\phi_{n_1,...,n_M}(\theta_1, ..., \theta_M) = \sqrt{\frac{\prod_{i=1}^{M} P_i(\theta_i)}{P(\theta_1, ..., \theta_M)}} \prod_{j=1}^{M} \phi_{n_j}(\theta_j), \tag{23}$$

given that the fraction $\prod_{i=1}^{M} P_i(\theta_i)/P(\theta_1, ..., \theta_M)$ is well behaved. The functions $\phi_{n_j}(\theta)$ are orthonormal with respect to the marginal distribution $P_j(\theta)$, i.e.:

$$\int P_j(\theta) \phi_{n_j}(\theta) \phi_{m_j}(\theta) d\theta = \delta_{n_j, m_j}. \tag{24}$$

The integration is done over the support of $P_i$. From this definition it immediately follows:

$$\int P(\theta_1, ..., \theta_M)\phi_{n_1,...,n_M}(\theta_1, ..., \theta_M)\phi_{m_1,...,m_M}(\theta_1, ..., \theta_M)d\theta_1 \cdots d\theta_M = \prod_{k=1}^{M} \delta_{n_k,m_k}. \tag{25}$$

The generalisation of Eq. 7 in S1 reads:

$$\hat{B}_{n_j,m_j} = \int \phi_{n_j}(\theta)\, \theta\, \phi_{m_j}(\theta)\, P_j(\theta)\, d\theta. \tag{26}$$

We denote by $\lambda^{(l_j)}$ the $l_j$th eigenvector of the matrix $\hat{B}$ and by $u_n^{(l_j)}$ the corresponding $n$th-component of the $l_j$th eigenvector. Applying an orthonormal basis transformation in a way similar to Eq. 22 in S1 yields:

$$\psi^s_{l_1,...,l_M}(\theta_1, ..., \theta_M) = \sum_{n_1=0}^{N} \cdots \sum_{n_M=0}^{N} u^{(l_1)}_{n_1+1} \cdots u^{(l_M)}_{n_M+1} \phi_{n_1,...,n_M}(\theta_1, ..., \theta_M) \tag{27}$$

Substituting Eq.23 in Eq. 27 gives rise to:

$$\psi^s_{l_1,...,l_M}(\theta_1, ..., \theta_M) = \sqrt{\frac{\prod_{i=1}^{M} P_i(\theta_i)}{P(\theta_1, ..., \theta_M)}} \prod_{j=1}^{M} \psi^s_{l_j}(\theta_j). \tag{28}$$

We further define:

$$\omega_{l_1,...,l_M} = \int P(\theta_1, ..., \theta_M)\psi^s_{l_1,...,l_M}(\theta_1, ..., \theta_M)d\theta_1 \cdots d\theta_M \tag{29}$$

using Eq. 28 yields:

$$\omega_{l_1,...,l_M} = \int \sqrt{\prod_{i=1}^{M} P_i(\theta_i) P(\theta_1, ..., \theta_M)} \prod_{j=1}^{M} \psi^s_{l_j}(\theta_j) d\theta_1 \cdots d\theta_M. \tag{30}$$

Finally, the expansion of a function $Y$ is given by:

$$Y^s(\theta_1, ..., \theta_M) = \sqrt{\frac{\prod_{i=1}^{M} P_i(\theta_i)}{P(\theta_1, ..., \theta_M)}} \sum_{l_1} \cdots \sum_{l_M} Y(\lambda^{(l_1)}, ..., \lambda^{(l_M)})\omega_{l_1,...,l_M} \psi^s_{l_1}(\theta_1) \cdots \psi^s_{l_M}(\theta_M). \tag{31}$$

In the case of uncorrelated parameters we recover Eq. 10 of the main text, because for $P(\theta_1, ..., \theta_M) = \prod_{i=1}^{M} P_i(\theta_i)$ we have $\omega_{l_1,...,l_M} = \prod_{i=1}^{M} u_1^{(l_i)}$.

#### 1.3 S3: Derivation of the probability density function for the exponential decay model

For the exponential decay model an analytical expression for the probability density function (PDF) can be obtained. In order to derive the PDF we start with:

$$P(y,t) = \int_0^\infty \delta(y - A(t,\theta))P(\theta)d\theta.$$

For log-normal distributed $k$ we have for the PDF of the uncertainty parameter $\theta$ and $k(\theta)$:

$$\begin{aligned} P(\theta) &= \frac{1}{\sqrt{2\pi}} e^{-\theta^2/2} \\ k(\theta) &= \mu e^{\alpha\theta - \alpha^2/2}, \end{aligned}$$

with

$$\alpha = \sqrt{\ln\left(1 + \frac{\sigma^2}{\mu^2}\right)}.$$

The function $A(t,\theta)$ is given by:

$$A(t,\theta) = A_0 e^{-k(\theta)t}.$$

Defing $u = y - x$ we arrive at:

$$P(y,t) = \int_{u(0)}^{u(\infty)} \delta(u) \left|\frac{\partial A}{\partial \theta}\right|^{-1}_{\theta=\theta(u)} P(\theta(u))du.$$

Now, it holds:

$$\begin{aligned} \frac{\partial A}{\partial \theta} &= -\alpha_0 t e^{-kt}\alpha k \\ \theta(u) &= \frac{1}{\alpha} \ln\left[\frac{e^{a^2/2}}{\mu t} \ln \frac{y-u}{x_0}\right], \quad u < y. \end{aligned}$$

and further:

$$\begin{aligned} u(\infty) &= y - A(t,\infty) = y - A_0 e^{-k(\infty)t} \\ k(\infty) &= \infty \quad (\alpha > 0) \\ u(\infty) &= \begin{cases} y - A_0 & : t = 0 \\ y & : t > 0 \end{cases} \\ k(-\infty) &= 0 \\ u(0) &= y - A_0 \end{aligned}$$

Using these results we arrive at:

$$
\begin{aligned}
P(y,t) &= \int_{u(0)}^{u(\infty)} \delta(u) \frac{e^{k(u)t}}{x_0 t \alpha k(u)} \frac{e^{-\theta^2(u)/2}}{\sqrt{(2\pi)}} du \\
P(y,t) &= \frac{e^{k(u=0)t}}{\sqrt{2\pi} A_0 t \alpha k(u=0)} e^{-\theta^2(0)/2} \theta(y) \theta(A_0 - y).
\end{aligned}
$$

Inserting

$$
k(\theta(u=0)) = \frac{1}{t} \ln\left(\frac{A_0}{y}\right),
$$

we finally arrive at:

$$
P(y,t) = \frac{1}{\sqrt{2\pi}\alpha y \ln\left(\frac{A_0}{y}\right)} \exp\left[-\frac{1}{2\alpha^2}\left\{\ln\left(\frac{e^{\alpha^2/2}}{\mu t}\ln\left(\frac{A_0}{y}\right)\right)\right\}^2\right] \theta(y)\theta(A_0 - y),
$$

which is the distribution plotted in ???

#### 1.4 Trichome model for MC and PCE comparison

The model used in this section consists of 6 proteins: TTG1, GL1, GL3, TRY, CPC and ETC. We also model the complex between TTG1 and GL3, and GL1 and GL3, as AC1 and AC2, respectively. This leads to the following set of equations:

$$
\begin{aligned}
\partial_t [TTG1]_j &= \theta_1 - [TTG1]_j(\theta_2 + \theta_3 [GL3]_j) + \theta_2 \theta_4 \hat{L}[TTG1]_j && (32) \\
\partial_t [GL1]_j &= \theta_5 + \theta_6 [AC2]_j - [GL1]_j(\theta_7 + \theta_8 [GL3]_j) + \theta_7 \theta_{30} \hat{L}[GL1]_j && (33) \\
\partial_t [GL3]_j &= \theta_9 + \frac{\theta_{10}\theta_{11}[AC1]_j^2}{\theta_{11} + [AC1]_j^2} + \frac{\theta_{12}\theta_{13}[AC2]_j^2}{\theta_{13} + [AC2]_j^2} - \\
&\quad [GL3]_j(\theta_{14} + \theta_3 [TTG1]_j + \theta_8 [GL1]_j + \theta_{15}[TRY]_j + && (34) \\
&\quad \theta_{16}[CPC]_j + \theta_{17}[ETC]_j) + \theta_{14}\theta_{31}\hat{L}[GL3]_j && (35) \\
\partial_t [TRY]_j &= \theta_{18}[AC1]_j^2 - [TRY]_j(\theta_{19} + \theta_{15}[GL3]_j) + \\
&\quad \theta_{19}\theta_{20}\hat{L}[TRY]_j && (36) \\
\partial_t [CPC]_j &= \theta_{21}[AC2]_j^2 - [CPC]_j(\theta_{22} + \theta_{16}[GL3]_j) + \\
&\quad \theta_{22}\theta_{23}\hat{L}[CPC]_j && (37) \\
\partial_t [ETC]_j &= \theta_{24}[AC1]_j^2 + \theta_{25}[AC2]_j^2 - [ETC]_j(\theta_{26} - \theta_{17}[GL3]_j) + \\
&\quad \theta_{26}\theta_{27}\hat{L}[ETC]_j && (38) \\
\partial_t [AC1]_j &= \theta_3 [GL3]_j [TTG1]_j - \theta_{28}[AC1]_j && (39) \\
\partial_t [AC2]_j &= \theta_8 [GL3]_j [GL1]_j - \theta_{29}[AC2]_j && (40)
\end{aligned}
$$

where $\hat{L}$ indicates the coupling equation between cells, given by

$$\begin{aligned}\hat{L}[\chi]_{x,y} =& [\chi]_{y-1,x} + [\chi]_{y+1,x} + [\chi]_{y,x-1} + [\chi]_{y,x+1} \\ &+ [\chi]_{y+1,x-1} + [\chi]_{y-1,x+1} - 6[\chi]_{y,x}.\end{aligned} \tag{41}$$

for any species $\chi$ and cell at coordinates $(x, y)$.

The parameters $\theta_1$ to $\theta_5$ were distributed lognormally with given mean and variance indicated in Table 1. Other parameters were fixed according to Table 2.

Table 1: Distribution parameters of uncertainty parameters used in MC and PCE comparison.

| $\theta_n$ | $\mu$ | $\sigma^2$ |
|---|---|---|
| $\theta_1$ | 7.7840 | 0.3892 |
| $\theta_2$ | 0.4621 | 0.0231 |
| $\theta_3$ | 1.6533 | 0.0827 |
| $\theta_4$ | 0.7828 | 0.0391 |
| $\theta_5$ | 0.4573 | 0.0229 |

Table 2: **Overview of parameters in the model given in equations (32) - (40) and their value.**

| $\theta_n$ | Value |
|---|---|
| $\theta_6$ | 3.6936 |
| $\theta_7$ | 0.6624 |
| $\theta_8$ | 5.2004 |
| $\theta_9$ | 0.2656 |
| $\theta_{10}$ | 1.5810 |
| $\theta_{11}$ | 3.4336 |
| $\theta_{12}$ | 0.1547 |
| $\theta_{13}$ | 1.4459 |
| $\theta_{14}$ | 0.8884 |
| $\theta_{15}$ | 2.9732 |
| $\theta_{16}$ | 0.1124 |
| $\theta_{17}$ | 3.0911 |
| $\theta_{18}$ | 0.7015 |
| $\theta_{19}$ | 2.1670 |
| $\theta_{20}$ | 6.5933 |
| $\theta_{21}$ | 3.8429 |
| $\theta_{22}$ | 6.7122 |
| $\theta_{23}$ | 0.1394 |
| $\theta_{24}$ | 0.1635 |
| $\theta_{25}$ | 1.5696 |
| $\theta_{26}$ | 3.7109 |
| $\theta_{27}$ | 6.6554 |
| $\theta_{28}$ | 0.9085 |
| $\theta_{29}$ | 0.5668 |
| $\theta_{30}$ | 2.5747 |
| $\theta_{31}$ | 0.1215 |
